# Microtubule forces drive nuclear damage in *LMNA* cardiomyopathy

**DOI:** 10.1101/2024.02.10.579774

**Authors:** Daria Amiad Pavlov, Julie Heffler, Carmen Suay-Corredera, Mohammad Dehghany, Kaitlyn M. Shen, Noam Zuela-Sopilniak, Rani Randell, Keita Uchida, Rajan Jain, Vivek Shenoy, Jan Lammerding, Benjamin Prosser

**Affiliations:** Department of Physiology, Pennsylvania Muscle Institute, Perelman School of Medicine, University of Pennsylvania; Weill Institute for Cell and Molecular Biology & Meinig School of Biomedical Engineering, Cornell University; Department of Materials Science and Engineering, Center for Engineering Mechanobiology, University of Pennsylvania; Departments of Medicine and Cell and Developmental Biology, Penn Cardiovascular Institute, Penn Epigenetics Institute, Perelman School of Medicine, University of Pennsylvania

## Abstract

Nuclear homeostasis requires a balance of forces between the cytoskeleton and nucleus. Mutations in the *LMNA* gene, which encodes the nuclear envelope proteins lamin A/C, disrupt this balance by weakening the nuclear lamina. This results in nuclear damage in contractile tissues and ultimately muscle disease. Intriguingly, disrupting the LINC complex that connects the cytoskeleton to the nucleus has emerged as a promising strategy to ameliorate *LMNA*-associated cardiomyopathy. Yet how LINC complex disruption protects the cardiomyocyte nucleus remains unclear. To address this question, we developed an assay to quantify the coupling of cardiomyocyte contraction to nuclear deformation and interrogated its dependence on the nuclear lamina and LINC complex. We found that, surprisingly, the LINC complex was mostly dispensable for transferring contractile strain to the nucleus, and that increased nuclear strain in lamin A/C*-*deficient cardiomyocytes was not rescued by LINC complex disruption. Instead, LINC complex disruption eliminated the cage of microtubules encircling the nucleus. Disrupting microtubules was sufficient to prevent nuclear damage and rescue cardiac function induced by lamin A/C deficiency. We computationally simulated the stress fields surrounding cardiomyocyte nuclei and show how microtubule forces generate local vulnerabilities that damage lamin A/C-deficient nuclei. Our work pinpoints localized, microtubule-dependent force transmission to the nucleus as a pathological driver and therapeutic target for *LMNA-* cardiomyopathy.

**Graphical abstract:** 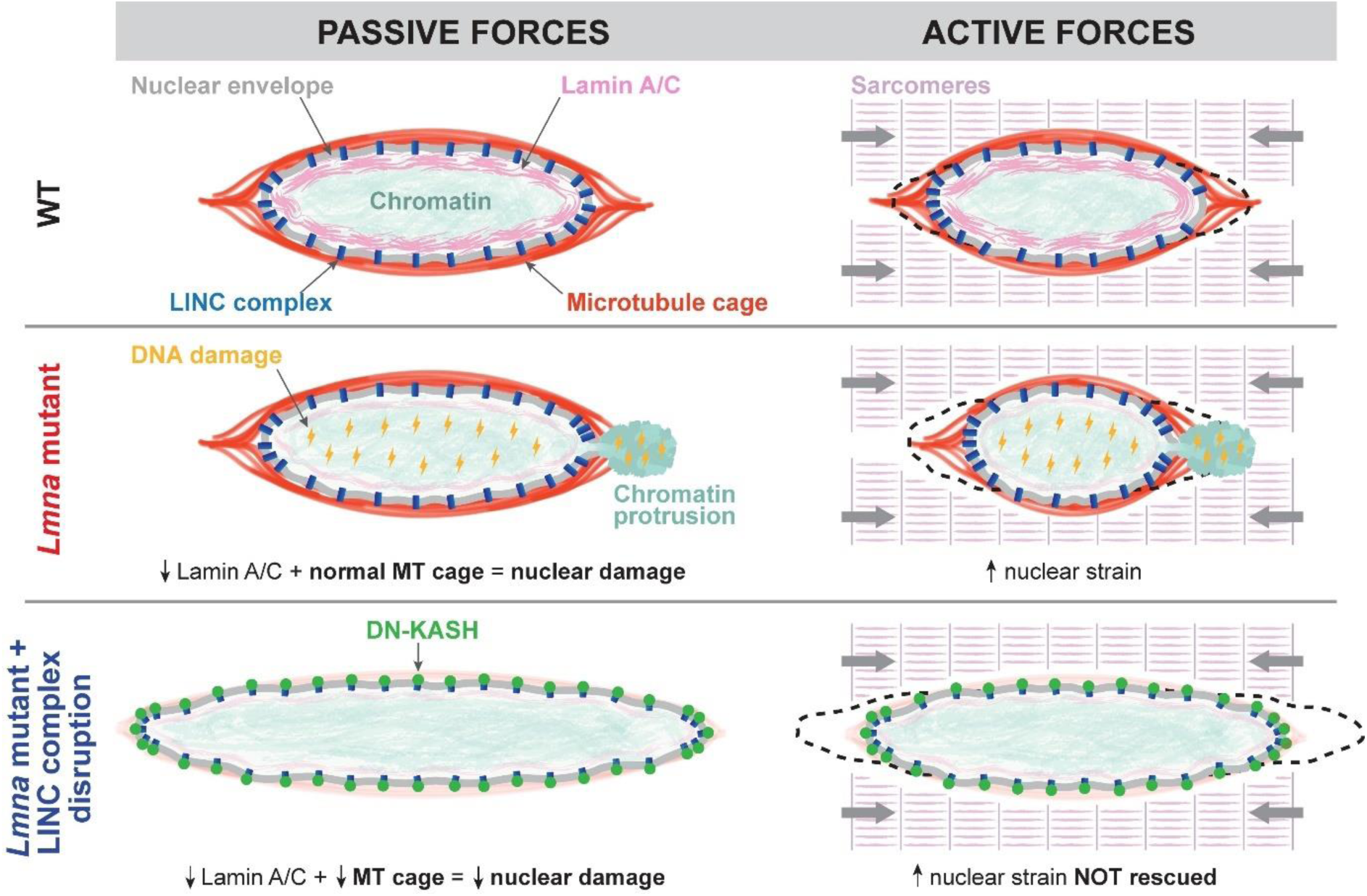

## Introduction

Cardiac muscle represents an extremely stressful mechanical environment with continuous contraction and relaxation cycles. Accordingly, the cytoskeleton and nucleus uniquely adapt to sense and withstand the mechanical load. In the adult cardiomyocyte, the actin-myosin, microtubule (MT), and desmin intermediate filament networks physically couple to the nucleus via the Linker of Nucleoskeleton and Cytoskeleton (LINC) complex that spans the nuclear envelope (NE). The LINC complex interacts directly with the nuclear lamina, a meshwork of A and B-type lamin filaments that provides structural support to the nucleus and contributes to spatial chromatin organization. The LINC complex consists of various Nuclear Envelope Spectrin Repeat (nesprin) protein isoforms at the outer nuclear membrane containing a conserved Klarsicht/ANC-1/Syne Homology (KASH) domain. The KASH domains interact across the luminal space with Sad1p/UNC-84 (SUN) domain-containing proteins SUN1 or SUN2 located at the inner nuclear membrane, which bind to the nuclear lamina and chromatin^1,2^. Thus, nuclear homeostasis and integrity depend on the balance of cytoskeletal forces, nuclear resistance, and their proper coupling through the LINC complex and nuclear lamina^3^.

The importance of this balance in the heart is demonstrated by variants in genes encoding lamins or other NE components that disproportionally affect highly contractile striated muscle tissues ^4,5^. Specifically, mutations in the *LMNA* gene that encodes A-type nuclear lamins (lamin A/C) cause a heterogeneous group of diseases (“laminopathies”) with high prevalence of cardiomyopathies ^6–8^. The *LMNA* N195K variant is associated with dilated cardiomyopathy in patients, and the *Lmna*^N195K/N195K^ mouse model leads to heart failure and death within 12 weeks of age ^9^. Specific and effective treatments are currently not available for *LMNA* cardiomyopathy, and the molecular mechanism of disease pathogenesis remains poorly understood. A prevailing hypothesis points to mechanically induced structural damage to the NE when the nuclear lamina is compromised ^10–15^.

Importantly, decoupling the nucleus from the cytoskeleton by disrupting the LINC complex restores cardiac function and extends the lifespan of *Lmna* cardiomyopathy mice ^16,17^. Yet the specific cytoskeletal forces responsible for nuclear damage in cardiomyocytes and the mechanism of protection by LINC complex disruption remain unclear ^10,13,17^. This is in part due to the lack of direct tools to measure how cytoskeletal forces are transferred to the nucleus in the relevant physiological and mechanical environment ^4,5,13^. Adult cardiomyocytes are often binucleated with nuclei equally spaced in the center of the cardiomyocyte, surrounded by a dense sarcomeric network and perinuclear cage of MTs. This central location prevents direct physical access to probe nuclear stiffness, as often reported in other cell types ^18^.

Here we introduce a new assay to quantify sarcomere-nuclear strain coupling in beating, adult cardiomyocytes. We use this assay to interrogate the role of the LINC complex and the cytoskeleton in driving nuclear strain during active cardiomyocyte contraction. We further investigate mechanisms of nuclear damage in *Lmna* cardiomyopathy and protection from nuclear damage by LINC complex disruption ^19^. Surprisingly, we found that sarcomere-nuclear strain coupling is largely preserved upon disruption of the LINC complex. Further, LINC complex disruption did not reduce contraction induced nuclear strain in *Lmna*-deficient cardiomyocytes. Instead, we observed that LINC complex disruption eliminates the dense cage of MTs normally formed around the nuclei of cardiomyocytes, which reduces MT-associated forces and confers protection from nuclear rupture at the nuclear tips. We found that MT disruption is sufficient to prevent nuclear damage in lamin A/C-deficient cardiomyocytes of mouse and human origin, and via a computational model that simulates the distribution of forces and sites of nuclear vulnerability in the cardiomyocyte. Finally, we demonstrate that in vivo disruption of the MT network is sufficient to preserve cardiac function and extend survival in mice following cardiomyocyte-specific *Lmna* depletion, further demonstrating the crucial role of MT-associated forces acting on the fragile nuclei in driving disease pathology in *Lmna* cardiomyopathy.

## Results

### Sarcomere-nuclear strain coupling in beating, adult cardiomyocytes

Measuring active mechanical signal transfer into the nucleus of mature cardiomyocytes is a missing component of interrogations into cardiac mechanobiology and *LMNA* cardiomyopathy. We developed a method to quantify sarcomere-nuclear strain coupling during electrically stimulated contractions in adult cardiomyocytes (Supplementary movie S1). We combined real-time measurements of sarcomere length change proximal to the nucleus with high spatiotemporal resolution (90 frames per second) Airyscan 2D imaging of the nucleus during steady-state (1 Hz) contraction (Fig. 1A). Figure 1B demonstrates that during the contraction cycle, sarcomere shortening and relaxation (magenta) are tightly coupled to the shortening and relaxation of nuclear length (cyan), with an opposing increase in nuclear width (orange). Notably, the relative change in sarcomere length (strain) during contraction is not fully transferred to the nucleus, resulting in dampening of nuclear strain (Fig. 1C). For example, at peak systole, for 11.4 ± 0.4 % sarcomere compression, nuclear length decreases only 6.6 ± 0.3 % (Fig. 1D). We generated sarcomere-nuclear strain coupling maps by plotting absolute sarcomere length versus absolute nuclear length during systolic compression and diastolic re-lengthening (Fig. 1E), or the respective sarcomere strain versus nuclear strain (Fig. 1F; unless otherwise specified, in our notation ‘strain’ refers to strain along the contractile axis). The dotted line represents a theoretical scenario where sarcomere strain would be linearly coupled, without loss, to nuclear strain. The upward deviation of the curve from the dotted line indicates that during contraction there is dampening, and not linear elastic coupling, between the sarcomere and nucleus. We can quantify this sarcomere-nuclear strain dampening during both phases of the contractile cycle (Fig. 1G). Systolic dampening is the integrated area under the curve during contraction with respect to the linear relationship (Fig. 1G, left). Diastolic dampening is the integrated area under the curve during re-lengthening with respect to a linear relationship with the intercept fixed at end-systolic strain (Fig. 1G, right). Using this approach, we can dissect how specific perturbations might compromise coupling between the sarcomeres and nucleus during each phase of the contractile cycle.

**Fig. 1:**
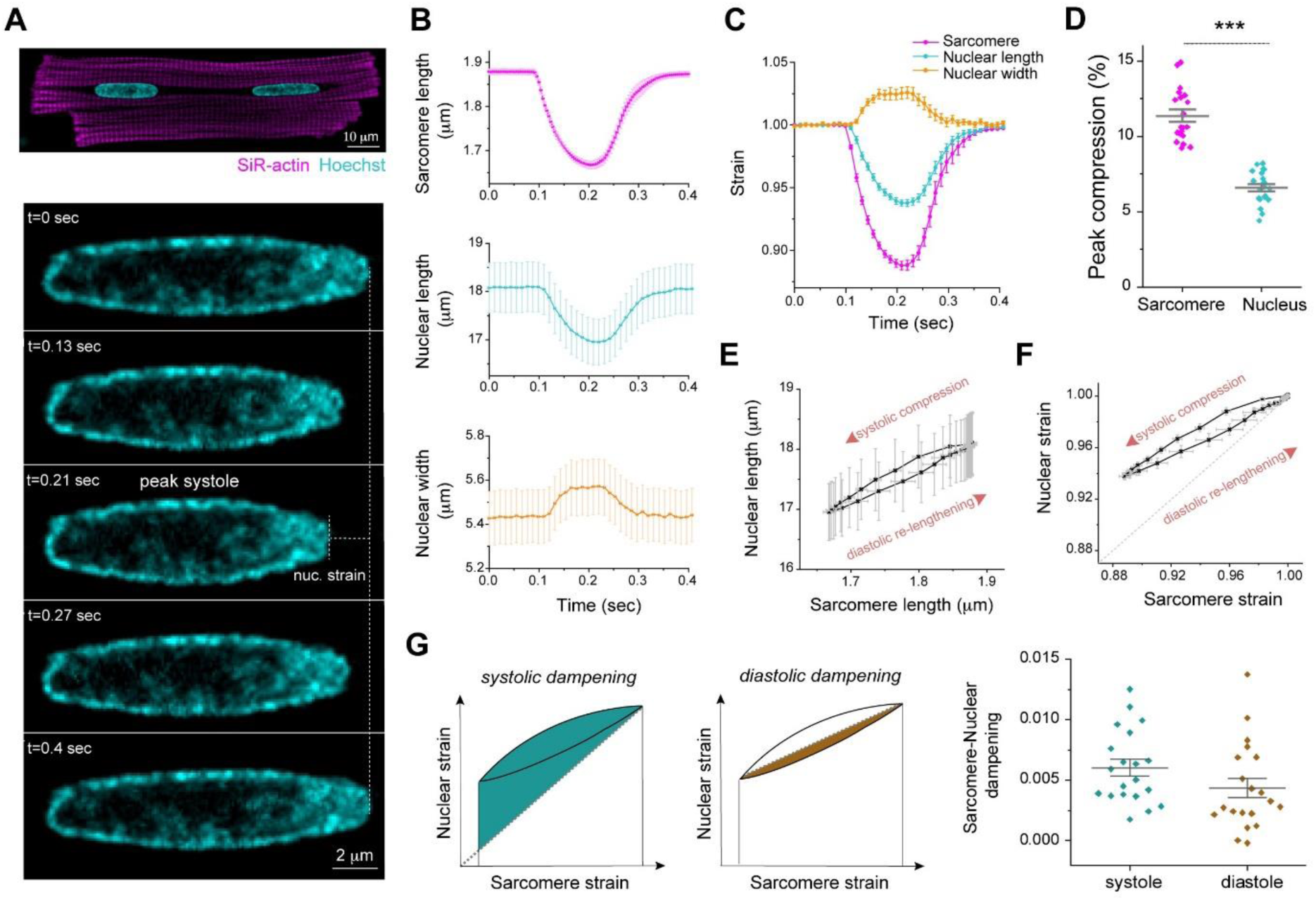
Active sarcomere-nuclear strain coupling in beating cardiomyocytes. (A) Live isolated adult rat cardiomyocyte stained with SiR-actin and Hoechst 33342 to visualize sarcomeres and DNA, respectively (top). Cardiomyocytes were stimulated at 1 Hz, and the nucleus imaged at 90 fps to follow nuclear deformation during the contraction cycle (bottom – representative time lapse images, see Supplementary movie S1). (B) Sarcomere length, nuclear length, and nuclear width recordings over time for a single contraction cycle. (C) Sarcomere strain, nuclear length strain and nuclear width strain over time. (D) Quantification of peak sarcomere and nuclear compression. (E) Sarcomere-nuclear coupling represented with a plot of nuclear length versus sarcomere length and (F) respective nucleus strain versus sarcomere strain. The latter, dimensionless strain coupling map depicts the dampened strain on the nucleus during systolic compression and diastolic re-lengthening, as marked by the deviation from a linear correlation (dotted line). (G) Sarcomere-nuclear strain dampening during systole quantified from area above linear correlation (left schematic). Diastolic dampening is quantified from area under end systolic linear correlation (right schematic). Data presented as mean ± standard error (SE) for 20 cells from a single representative adult rat heart. Statistical significance determined by two-tailed t-test (***, *p* < 0.001).

We next interrogated how cytoskeletal connections to the nucleus regulate sarcomere-nuclear strain coupling during contraction. To disrupt all cytoskeletal interactions with the LINC complex, we utilized a previously validated adenovirus over-expressing a dominant-negative KASH peptide (AdV DN-KASH), which prevents interactions between endogenous nesprin and SUN proteins ^1,3^. To disrupt MTs, we used colchicine (colch, 1 µM, overnight) (Fig. 2A). After 48 h AdV transduction of isolated rat cardiomyocytes, we confirmed that AdV DN-KASH disrupted the LINC complex through loss of perinuclear endogenous nesprin 1 (Extended Data Fig. 1A), and that 24 h colchicine treatment resulted in loss of MTs (Extended Data Fig. 1B). Neither acute LINC complex disruption nor MT depolymerization overtly altered sarcomere organization around the nucleus (Extended Data Fig. 1A-B) or resting sarcomere length (Extended Data Fig. 1C).

**Fig. 2:**
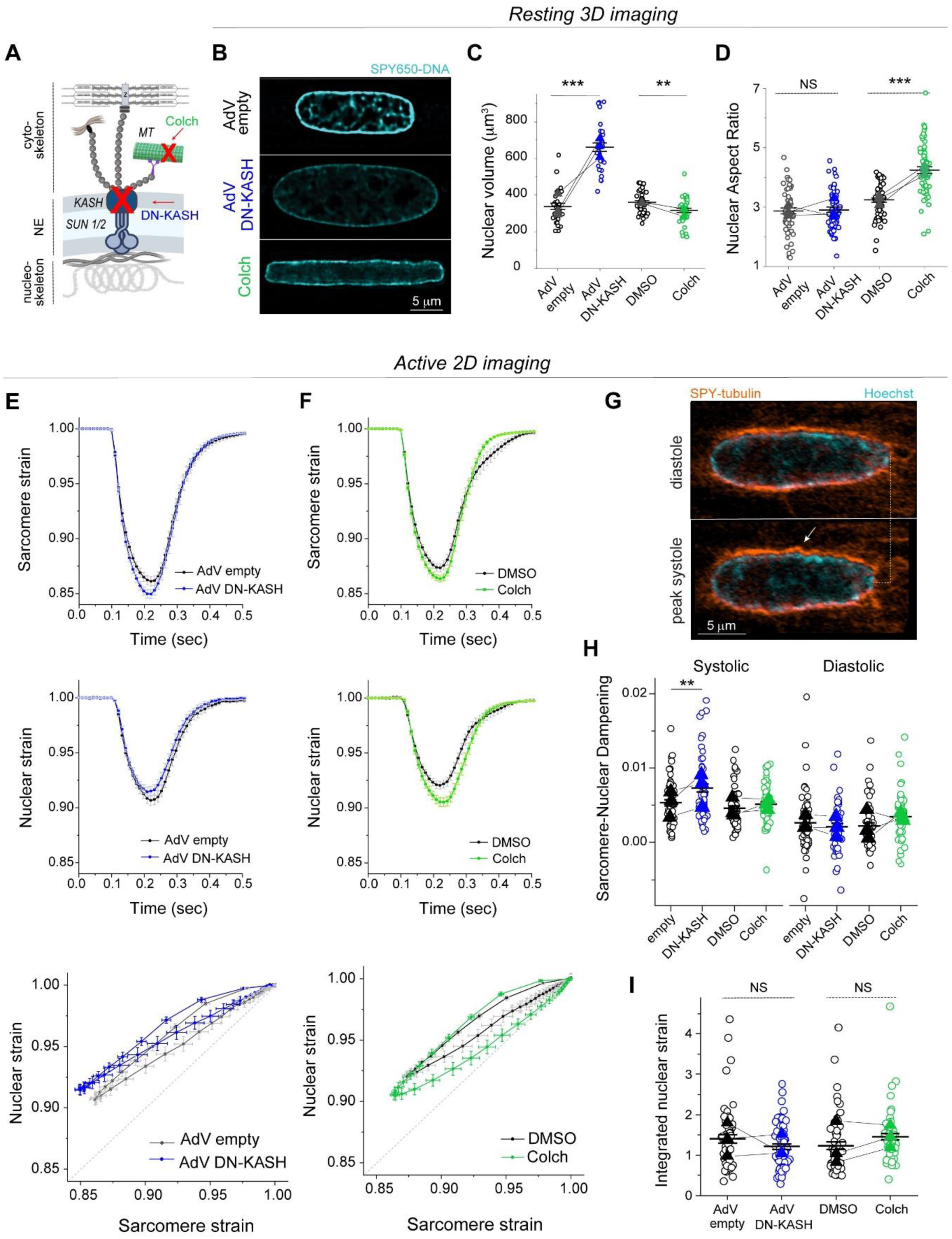
**Distinct effects of LINC complex disruption and MT depolymerization on resting nuclear morphology and active sarcomere-nuclear strain coupling**. (A) Schematic of cytoskeletal to nucleoskeletal connections and the experimental perturbations. (B) Live 3D super-resolution imaging of resting adult rat cardiomyocyte nuclei (mid-nuclear planes are displayed). Nuclear volume (C) and aspect ratio (D) measurements following AdV DN-KASH or colchicine treatment. For panel (C): AdV empty and AdV DN-KASH (48 h): *N* = 3, *n* = 30, DMSO and colch (24 h): *N* = 3, *n* = 36. For panel (D): AdV empty and AdV DN-KASH (48 h): *N* = 3, *n* = 51, DMSO and colch (24h): *N* = 3, *n* = 51. (E-I) Active 2D imaging of electrically stimulated cardiomyocytes. Sarcomere strain (top) and nuclear strain (middle) over time, and sarcomere-nuclear stain coupling (bottom) for (E) AdV DN-KASH and (F) colchicine compared to their respective controls. (G) Snapshots of live, WT cardiomyocyte labeled with SPY-555 tubulin and Hoechst 33342 during diastole and peak systole demonstrating MT cage buckling (arrow) during contraction. (H) Quantification of sarcomere-nuclear strain dampening during the systolic and diastolic phases, for the indicated perturbations (see schematics in Fig. 1G). (I) Integrated nuclear strain over time during the contractile cycle for the indicated perturbations. AdV empty and AdV DN-KASH (48 h): *N* = 3, *n* = 51, DMSO and colch (24 h): *N* = 3, *n* = 51. Data presented as mean ± SE on individual cells. For panels C, D, H, I individual cells are indicated with open circles and individual animal replicate means are indicated with closed triangles connected by lines between experimental groups. Statistical significance determined by two-tailed t-test (*, *p* < 0.05; **, *p* < 0.01; ***, *p* < 0.001).

We first assessed the effect of acute LINC complex and MT disruption on nuclear morphology via live, 3D super-resolution confocal imaging (AiryScan jDCV) of quiescent (non-contracting) adult rat cardiomyocytes (Fig. 2B). AdV DN-KASH increased nuclear size in all 3 dimensions, leading to a substantial increase in nuclear volume (Fig. 2C). Colchicine treated nuclei displayed elongated morphology with increased nuclear aspect ratio and a subtle decrease in nuclear volume (Fig. 2C-D), in agreement with our earlier report and consistent with a reduction of compression on the nucleus ^3^.

We then assessed the active sarcomere-nuclear strain coupling via rapid 2D imaging in electrically stimulated (1 Hz) and contracting cardiomyocytes. Upon electrical excitation, peak sarcomere contractility increased subtly with AdV DN-KASH, yet peak nuclear compression *decreased* slightly (Fig. 2E, top and middle, Extended Data Fig. 1C, left). This indicated a modest decrease in sarcomere-nuclear strain coupling during myocyte contraction, as evident from the upward shift of the AdV DN-KASH curve in the strain coupling map (Fig. 2E bottom), and the significantly increased sarcomere-nuclear dampening during systole (Fig. 2H).

Colchicine treatment moderately increased sarcomere contraction and accelerated sarcomere relaxation (Fig. 2F, top), consistent with MTs normally providing a viscoelastic resistance to sarcomere motion ^20^. Nuclear compression proportionately increased, but nuclear relaxation did not accelerate to match the faster sarcomere relaxation (Fig. 2F, top and middle). Faster sarcomere vs. nuclear re-lengthening manifests as increased area, or hysteresis, within the diastolic strain coupling curve upon colchicine treatment (Fig. 2F, bottom). Buckling of the MT cage during systole (Fig. 2G, Supplementary movie S2) demonstrates the transmission of contractile forces to the nucleus through the MT cage. This buckling might provide restoring force to match sarcomere and nuclear relaxation rates during diastole ^21^. To examine the cumulative strain on the nucleus during the full contractile cycle, we integrated the nuclear strain over time, and found no statistically significant differences between control cells and cells with LINC complex disruption or colchicine induced MT disruption (Fig. 2I). Together, these results demonstrate that acute, in vitro LINC complex disruption subtly reduces sarcomere-nuclear strain coupling, but neither LINC complex nor MT disruption significantly alter integrated nuclear strain during the contractile cycle. This indicates that nuclear strain during cardiomyocyte contraction is driven primarily by the shortening and re-lengthening of nearby sarcomeres, independent of their connectivity to the nucleus via the LINC complex.

### Cardiac-specific, in vivo LINC complex disruption protects from nuclear damage and cardiac dysfunction in *Lmna* N195K mice

We next sought to investigate the role of cytoskeletal forces in nuclear damage driven by lamin A/C deficiency or disease-causing *Lmna* mutations. We utilized the previously described *Lmna*^N195K/N195K^ (hereafter ‘*Lmna* N195K*’*) mutant mouse model that exhibits heart failure and death by 12 weeks of age ^9^. The *Lmna* N195K mutation leads to mechanical fragility of the nucleus, with a loss in nuclear stability similar to that observed with complete deletion of lamin A/C ^10,11^. Recent studies found that global and cardiac-specific LINC complex disruption improves cardiac function and prolongs lifespan in *Lmna* N195K and other *Lmna* mutant mice ^16,17^. To probe the cardiomyocyte-specific effects of LINC complex disruption, we used the previously described inducible *αMHC MerCreMer* DN-KASH mouse model, in which a dominant negative KASH domain of nesprin-2 tagged with GFP is expressed in cardiomyocytes upon tamoxifen treatment (referred to as cardiac-specific DN KASH, or csDN-KASH)^19^. We crossed these mice to *Lmna* N195K mice to generate a *Lmna* cardiomyopathy model with cardiac-specific inducible LINC complex disruption. Fig. 3A summarizes the different mouse models used in the next several figures. LINC complex disruption was induced by 5 consecutive daily tamoxifen injections at 3-4 weeks of age, followed by cardiomyocyte isolation at 8-9 weeks of age, allowing for 5 weeks of in vivo LINC complex disruption. We confirmed successful csDN-KASH induction following tamoxifen treatment by scoring the appearance of a csDN-KASH-GFP ring that surrounded the nucleus in >95% of cardiomyocytes, consistent with previous estimates of induction efficiency with this model ^19^. We confirmed LINC complex disruption upon csDN-KASH induction in cardiomyocytes by the loss of perinuclear nesprin-1 and nesprin-2 (Extended Data Fig. 2A-B). Consistent with previous reports ^16,17^, LINC complex disruption significantly extended the lifespan of *Lmna* N195K mice (Fig. 3B), improved cardiac contractility (Fig. 3C) and structure (Extended Data Fig. 2D), and reduced fibrosis as quantified from cardiac tissue slices (Fig. 3D). Beyond the 10-week time point, cardiac function was largely maintained in *Lmna* N195K csDN-KASH mice even at 12 weeks of age, when most of the *Lmna* N195K controls without LINC complex disruption had already died (Extended Data Fig. 2C).

**Fig. 3:**
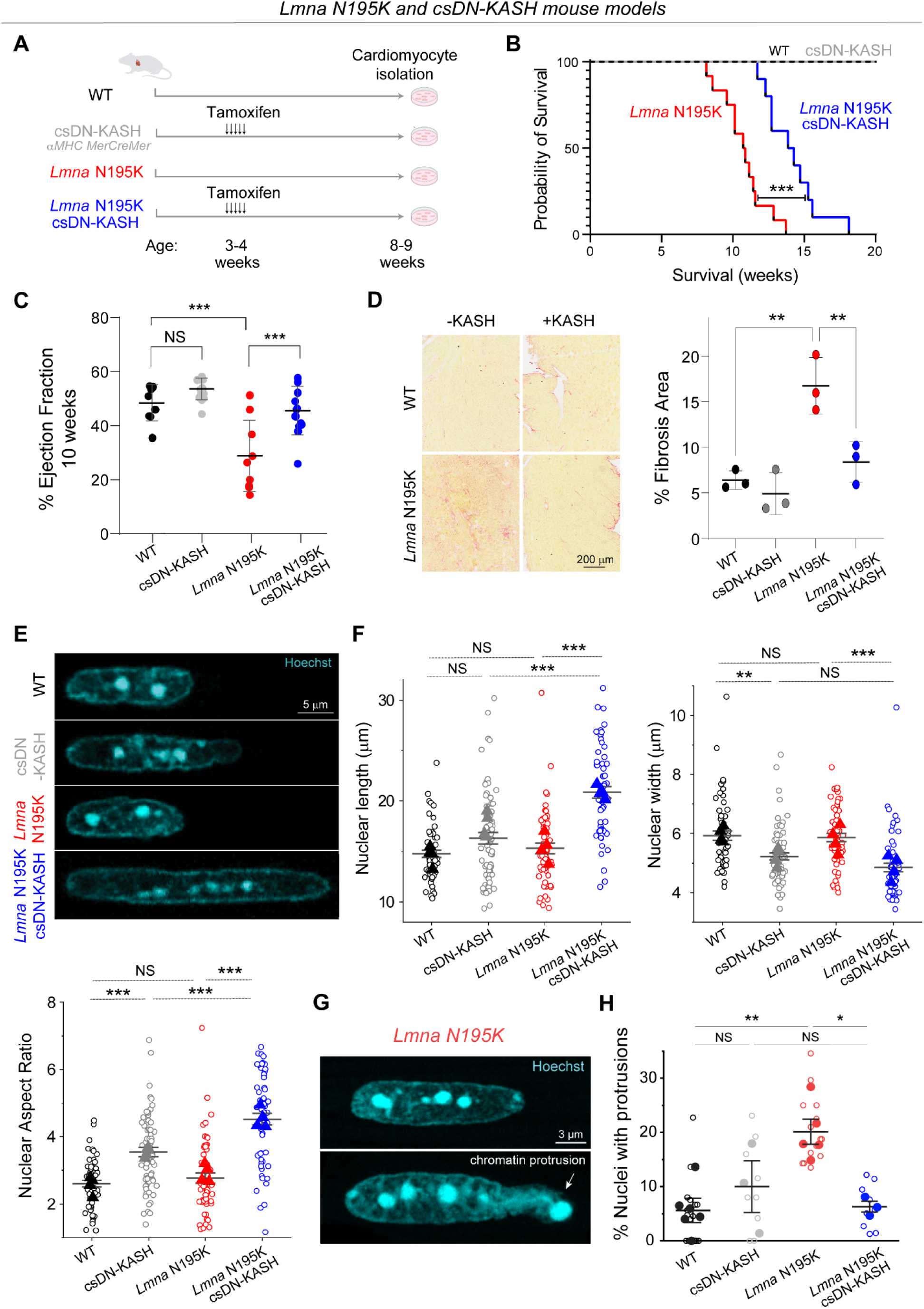
Cardiac specific LINC complex disruption extends lifespan, improves cardiac function and protects against nuclear ruptures in *Lmna* N195K cardiomyopathy. (A) Schematic presentation of the mouse models and experimental timelines used in this study. (B) Kaplan-Meier survival plot of the different experimental groups. N=10 mice per genotype. (C) Left ventricular ejection fraction measured by echocardiography, at 10 weeks of age. WT: *N* = 9, csDN-KASH: *N* = 15, *Lmna* N195K: *N* = 9, *Lmna* N195K csDN-KASH: *N* = 13. Error bar represents mean ± SD. Statistical significance determined by one-way ANOVA with Tukey multiple correction. (D) Representative Picrosirius red--stained heart sections and quantification of % fibrotic area for the indicated groups. *N* = 3 animal replicates per group. Error bar represents mean ± SD. Statistical significance determined by one-way ANOVA with Tukey multiple correction. (E) Representative images of cardiomyocyte nuclear morphology at 8-9 weeks of age for the studied groups. (F) Quantification of nuclear length, width, and aspect ratio. WT: *N* = 4, *n* = 58. csDN-KASH: *N* = 4, *n* = 69. *Lmna* N195K: *N* = 4, *n* = 55. *Lmna* N195K csDN-KASH: *N* = 4, *n* = 59. Data presented as mean ± SE of individual nuclei (open circles) while individual animal replicate means are indicated with closed triangles. (G) Chromatin protrusion from the nucleus in a *Lmna* N195K cardiomyocyte, suggestive of nuclear rupture. Representative mid plane images of a normal *Lmna* N195K nucleus (top) and a nucleus with a chromatin protrusion (bottom) in *Lmna* N195K cardiomyocytes. (H) Quantification of the percentage of nuclei with chromatin protrusions in each genotype by three independent, blinded scorers. Open circles represent the percentage of nuclei with chromatin protrusions quantified by each independent user for each animal. Closed circles correspond to the mean for each animal. Mean ± SE is shown for each genotype. WT: *N* = 5, *n* = 223. csDN-KASH: *N* = 3, *n* = 75. *Lmna* N195K: *N* = 5, *n* = 287. *Lmna* N195K csDN-KASH: *N* = 3, *n* = 223. Statistical significance determined by 1-way ANOVA with Bonferroni correction (*, *p* < 0.05; **, *p* < 0.01; ***, *p* < 0.001).

We next assessed resting nuclear morphology using high-resolution imaging of isolated cardiomyocytes from the four mouse groups. We did not observe significant changes in nuclear dimensions between WT and *Lmna* N195K cardiomyocytes with this approach (Fig. 3E-F). A separate analysis of a larger, albeit lower resolution data set revealed a subtle (11%) but statistically significant increase in nuclear aspect ratio in *Lmna* N195K cardiomyocytes (Extended Data Fig. 3A).

In contrast, LINC complex disruption (csDN-KASH) resulted in substantial elongation of WT cardiomyocyte nuclei (36% increase in nuclear aspect ratio) and even more extreme nuclear elongation in *Lmna* N195K mice (*Lmna* N195K csDN-KASH), which exhibited a 63% increase in nuclear aspect ratio (Fig. 3E-F). This nuclear elongation occurred independent of any corresponding changes in cellular length, width, or aspect ratio (Extended Data Fig. 3B), instead indicating an altered balance between cytoskeletal forces acting on the nucleus and internal nuclear resistance ^3^. Notably, the increased nuclear aspect ratio due to in vivo LINC complex disruption mirrored the alterations in nuclear morphology upon colchicine treatment (Fig. 2D, Fig. 3F), suggesting it may arise from a reduction in MT compressive forces.

In addition, we observed increased nuclear fragility and NE rupture in *Lmna* N195K vs. WT nuclei, scored as chromatin that appeared to spill out of the nucleus, with the nuclear lamina only partially enclosing the protrusion (Fig. 3G, Extended data Fig 3C). In agreement with a recent report of nuclear ruptures in cardiac specific *Lmna* KO mice ^15^, we observed chromatin protrusions in mature cardiomyocytes exclusively at the tips of nuclei. Using blinded scoring, we identified chromatin protrusions in 20% of nuclei from *Lmna* N195K cardiomyocytes (Fig. 3H, see Methods for details on scoring criteria), similar to recent results reported in lamin A/C-depleted cardiomyocytes ^15^. Importantly, in vivo LINC complex disruption rescued NE rupture events in *Lmna* N195K mice back to levels of WT controls (Fig. 3H). Taken together, these findings confirm the protective effect of LINC complex disruption on lifespan and cardiac function in *Lmna* N195K mice, and further demonstrate that LINC complex disruption leads to a marked protection from NE ruptures.

### In vivo LINC complex disruption does not reduce nuclear strain during myocyte contraction

A prevailing hypothesis for NE ruptures in laminopathies is that fragile nuclei are more susceptible to mechanical damage induced by forceful sarcomeric contractions ^13^. Therefore, decoupling the nucleus from the cytoskeleton might reduce contractility-induced nuclear damage ^16^. To probe this hypothesis, we first asked whether *Lmna* N195K cardiomyocyte nuclei indeed undergo more strain during the contractile cycle. Representative images of *Lmna* N195K nuclei at diastole and peak systole are depicted in Fig. 4A. Consistent with a more deformable nucleus ^10^, *Lmna* N195K cardiomyocytes showed increased nuclear compression during sarcomere contraction (Extended Data Fig. 4A) and nuclear re-lengthening that lagged behind sarcomere re-lengthening. This resulted in a downward shift in the strain coupling curve (Fig. 4B), increased diastolic sarcomere-nuclear dampening (Fig. 4G), and a significant increase in integrated nuclear strain over the contractile cycle in *Lmna* N195K cardiomyocytes (Fig. 4H). These data provide the first direct evidence for increased nuclear strain upon contraction of mutant *Lmna* mature cardiomyocytes.

**Fig. 4:**
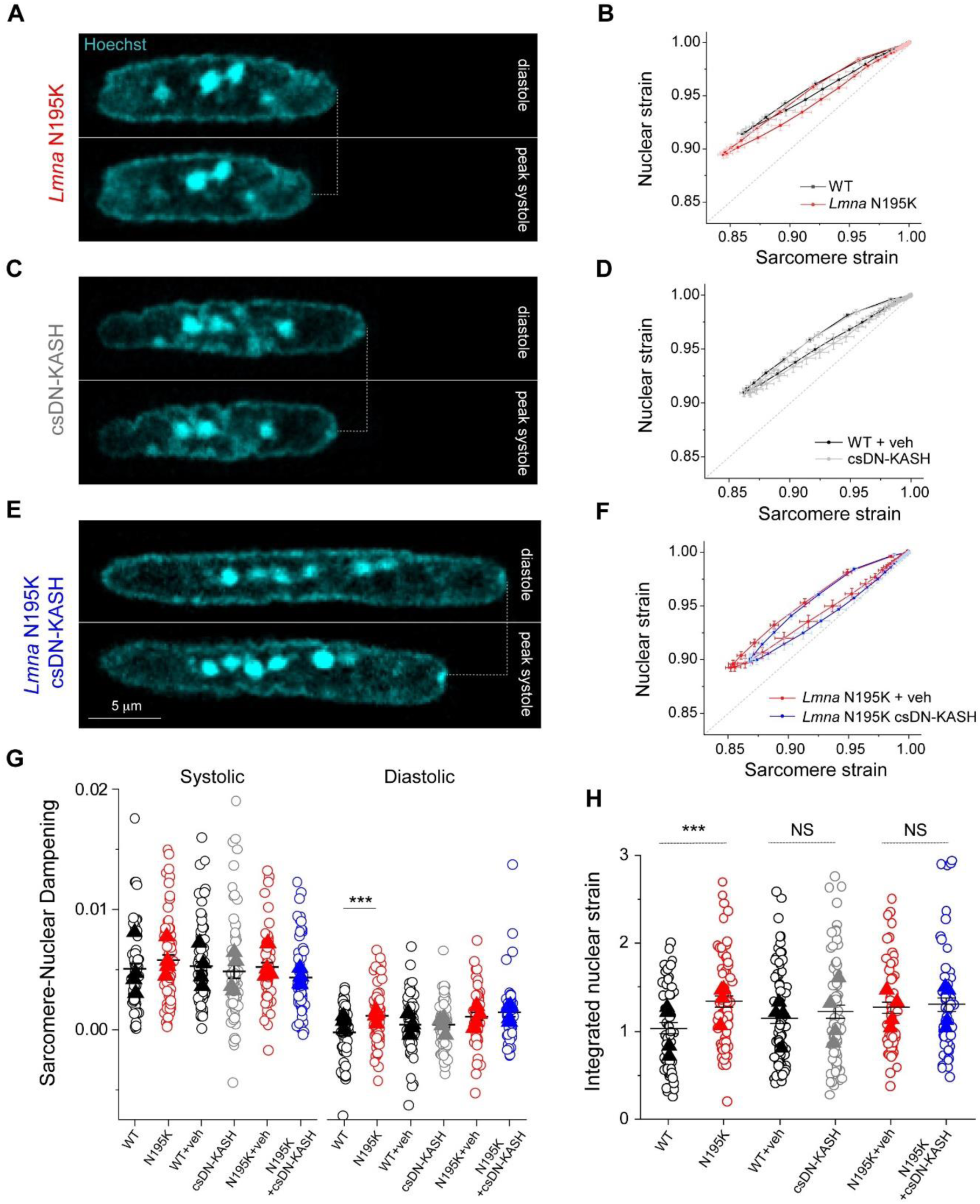
**Increased active nuclear strain in *Lmna* cardiomyopathy is not restored by cardiac specific in vivo LINC complex disruption**. (A) Representative snapshots of *Lmna* N195K cardiomyocyte nuclei during diastole and peak systole. (B) Increased nuclear compression as evidenced by a downward shift in the laminopathy strain coupling curve. (C) Representative snapshots of cardiac specific LINC complex disruption in WT cardiomyocyte nuclei during diastole and peak systole with (D) no change in active strain coupling. (E) Representative snapshots of cardiac specific LINC complex disruption in *Lmna* N195K cardiomyocyte nuclei during diastole and peak systole with (F) no change in active strain coupling. Data presented as mean ± SE (G) Quantification of sarcomere-nuclear strain dampening during the systolic and diastolic phases, for the indicated groups (see schematics in Fig. 1G). (H) Integrated nuclear strain over time during the contractile cycle for the indicated groups. WT: *N* = 4, *n* = 58. *Lmna* N195K: *N* = 4, *n* = 69. WT veh: *N* = 4, *n* = 81. csDN-KASH: *N* = 4, *n* = 69. *Lmna* N195K veh: *N* = 4, *n* = 55. *Lmna* N195K csDN-KASH: *N* = 4, *n* = 59. Data presented as mean ± SE of individual cells (open circles) while individual animal replicate means are indicated with closed triangles. Statistical significance determined by two-tailed t-test (*, *p* < 0.05; **, *p* < 0.01; ***, *p* < 0.001).

We next hypothesized that LINC complex disruption reduces the increased nuclear strain in *Lmna* N195K cardiomyocytes. Representative images of nuclei from csDN-KASH and *Lmna* N195K csDN-KASH mice at diastole and peak systole are depicted in Fig. 4C and 4E, respectively. Surprisingly, nuclear compression was not reduced in csDN-KASH cardiomyocytes (Extended Data Fig. 4B), and the overall strain coupling curve (Fig. 4D) and sarcomere-nuclear dampening (Fig. 4G) remained unchanged. These data further indicate that an intact LINC complex is not required for transferring the majority of sarcomeric strain into the nucleus during myocyte contraction. Consistently, neither nuclear compression (Extended Data Fig. 4C), sarcomere-nuclear dampening (Fig. 4G), nor active strain coupling (Fig. 4F) were altered in *Lmna* N195K csDN-KASH myocytes compared to *Lmna* N195K controls. The increased integrated nuclear strain in *Lmna* N195K mutants was not rescued by in vivo LINC complex disruption (Fig.4H). These results indicate that although the more deformable *Lmna* N195K nuclei undergo increased strain during the contractile cycle, this effect is not rescued by LINC complex disruption, yet LINC complex disruption reduces nuclear damage and improves cardiac function and overall survival. These findings suggests that LINC complex disruption confers cardioprotection through an alternative mechanism, independent of reducing nuclear strain during cardiomyocyte contraction.

### In vivo LINC complex disruption eliminates the perinuclear microtubule cage

We next explored alternative sources of mechanical stress that could damage *Lmna* deficient nuclei and be reduced by LINC complex disruption. We hypothesized that nuclear elongation upon csDN-KASH might be a consequence of reduced MT compression on the nucleus. Consistently, we observed almost complete elimination of the perinuclear MT cage (measured as perinuclear to cytosolic α-tubulin enrichment) upon csDN-KASH induction in WT and *Lmna* N195K cardiomyocytes (Fig. 5A-B). The dense perinuclear MT cage in cardiomyocytes was specifically enriched at nuclear tips (Extended Data Fig. 5A), which might contribute to increased mechanical forces at this location. This appeared noteworthy given that chromatin protrusions in *Lmna* N195K cardiomyocytes were exclusively observed at nuclear tips (Fig. 3G, Extended data Fig 3C).

**Fig. 5:**
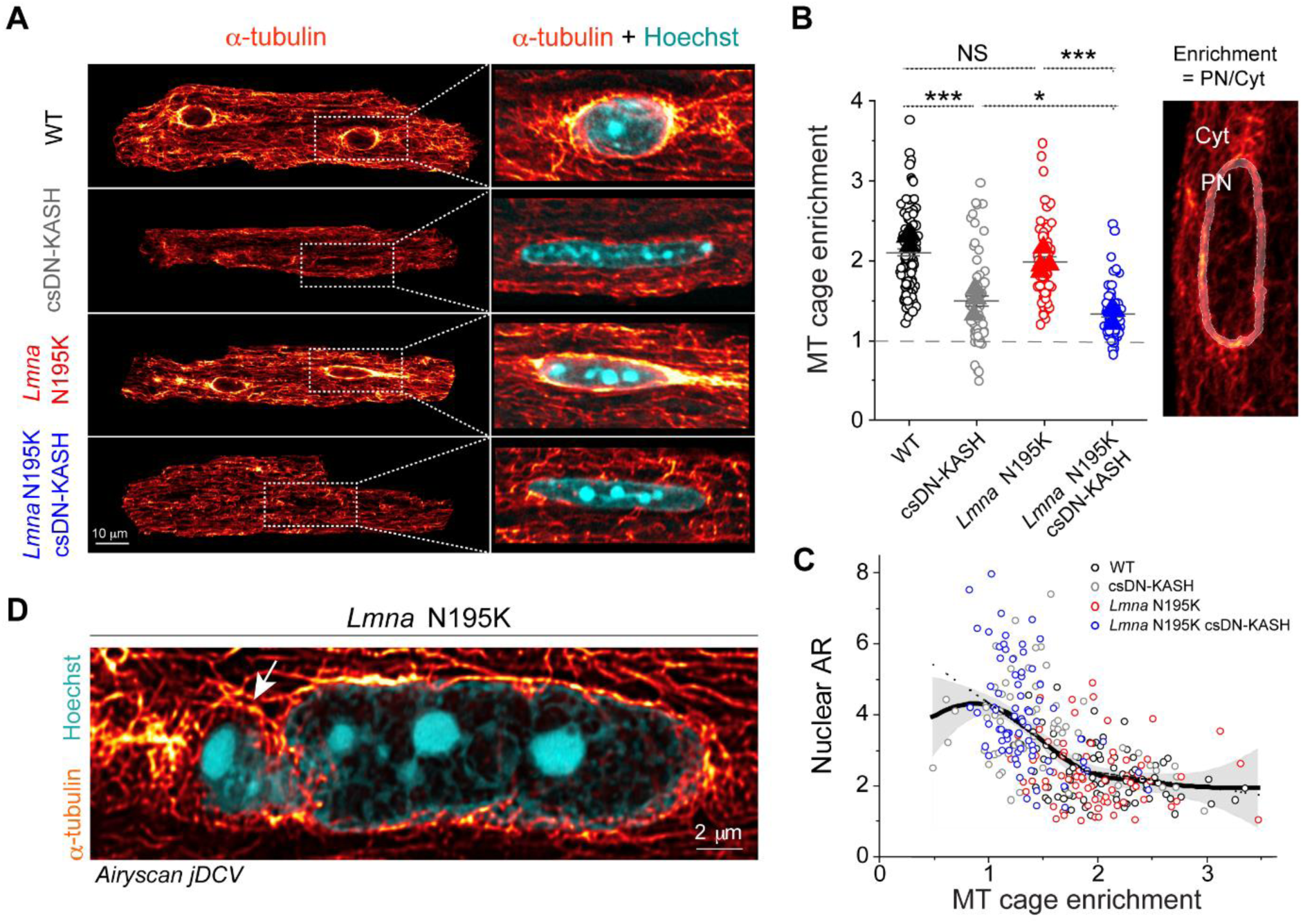
In vivo LINC complex disruption eliminates perinuclear MT cage leading to nuclear elongation. (A) Representative mid-plane immunofluorescence images of the MT network (α-tubulin), with a zoom in on a merge with labeled nuclei (Hoechst) in the WT, csDN-KASH, *Lmna* N195K, and *Lmna* N195K csDN-KASH adult mouse cardiomyocytes. (B) Quantification of perinuclear MT enrichment defined as perinuclear (PN) to cytoplasmic (Cyt) α-tubulin ratio (illustrated on the image on the right). Data presented as mean ± SE of individual nuclei (open circles) while individual animal replicate means are indicated with closed triangles. Statistical significance determined by one-way ANOVA with Bonferroni correction (*, *p* < 0.05; **, *p* < 0.01; ***, *p* < 0.001). (C) Nuclear aspect ratio (AR) as a function of perinuclear MT enrichment. Open circles represent individual nuclei from the respective groups. ‘Loess’ smoothing trace depicted with continuous black line and gray error area. Biphasic piecewise linear regression fit is indicated with a dashed black line. (D) Super resolution Airyscan jDCV mid plane image of a *Lmna* N195K cardiomyocyte nucleus (Hoechst) with the dense MT network (α-tubulin) penetrating the chromatin protrusion. WT: *N* = 3, *n* = 85. csDN-KASH: *N* = 3, *n* = 71. *Lmna* N195K: *N* = 3, *n* = 70. *Lmna* N195K csDN-KASH: *N* = 3, *n* = 75.

We leveraged intracellular heterogeneity to check if the nuclear aspect ratio depended on the levels of perinuclear MT enrichment. We observed a biphasic relationship between nuclear aspect ratio and the perinuclear MT cage (Fig. 5C). A ‘Loess’ smoothing algorithm applied to the pooled data from all experimental groups (Fig. 5C continuous black trace with gray error area) revealed a deflection point at ‘MT cage enrichment’ = 1.9. This deflection point was further verified by fitting a piecewise linear regression (dashed black trace) which indicated a negative correlation between nuclear aspect ratio and MT cage enrichment below 1.9 (slope = –2.1, *p* < 0.001). In contrast, the slope above this deflection point was flat (*p* > 0.05) indicating constant nuclear aspect ratio for MT cage enrichment ≥ 1.9. Together, these observations support the hypothesis that nuclear elongation upon in vivo LINC complex disruption is likely driven by decoupling of the MT network from the nucleus, with even greater elongation in more deformable *Lmna* N195K nuclei.

We thus hypothesized that nuclear damage in *Lmna* N195K nuclei is driven by perinuclear MT-dependent forces. In support of this hypothesis, super-resolution images of the MT network in *Lmna* N195K cardiomyocytes showed MTs encaging the protruded chromatin at the nuclear tip (Fig. 5D). Desmin intermediate filaments normally help balance compressive MT forces on the nucleus in mature cardiomyocytes ^3^; thus desmin deficiency could potentially contribute to such nuclear damage. However, we did not observe any decrease in the levels of desmin or alterations in the organization of the desmin network between the groups (Extended Data Fig. 5B-C). These data suggest that nuclear fragility, and not desmin deficiency, facilitates MT-based nuclear damage in the *Lmna* N195K cardiomyocytes.

MTs interact with the LINC complex via kinesin motors, and nesprins are required to recruit kinesin motors and MTs to the nuclear periphery ^22^. We found that LINC complex disruption depletes kinesin-1 from the perinuclear space (Extended Data Fig. 5D), in agreement with earlier reports ^10,17,23^. We also observed robust perinuclear kinesin-1 depletion with partial loss of the perinuclear MT cage following 48 h AdV DN-KASH transduction in isolated adult rat cardiomyocytes (Extended Data Fig. 5E-F), confirming our findings in an independent adult cardiomyocyte model. These findings suggest that LINC complex disruption leads to rapid loss of perinuclear kinesin motors and eventual loss of the perinuclear MT cage.

### *Lmna* N195K mutant and lamin A/C-deficient cardiomyocytes have increased NE ruptures linked to perinuclear MT network

To directly test the role of MTs in nuclear ruptures, we used transgenic mice expressing a fluorescent cGAS-tdTomato reporter ^10^ crossed with the *Lmna* N195K mouse model to allow quantification of NE rupture in live cardiomyocytes. Live imaging of freshly isolated cardiomyocytes revealed cGAS-tdTomato foci in and around the nucleus in the *Lmna* N195K group that were significantly increased compared to cGAS-tdTomato WT mice (Fig. 6A and Extended Data Fig. 6A). In *Lmna* N195K cardiomyocytes, cGAS foci were mostly observed at the nuclear tips, where MTs were enriched, regardless of the absence or presence of chromatin protrusions (Fig. 6B).

**Fig. 6:**
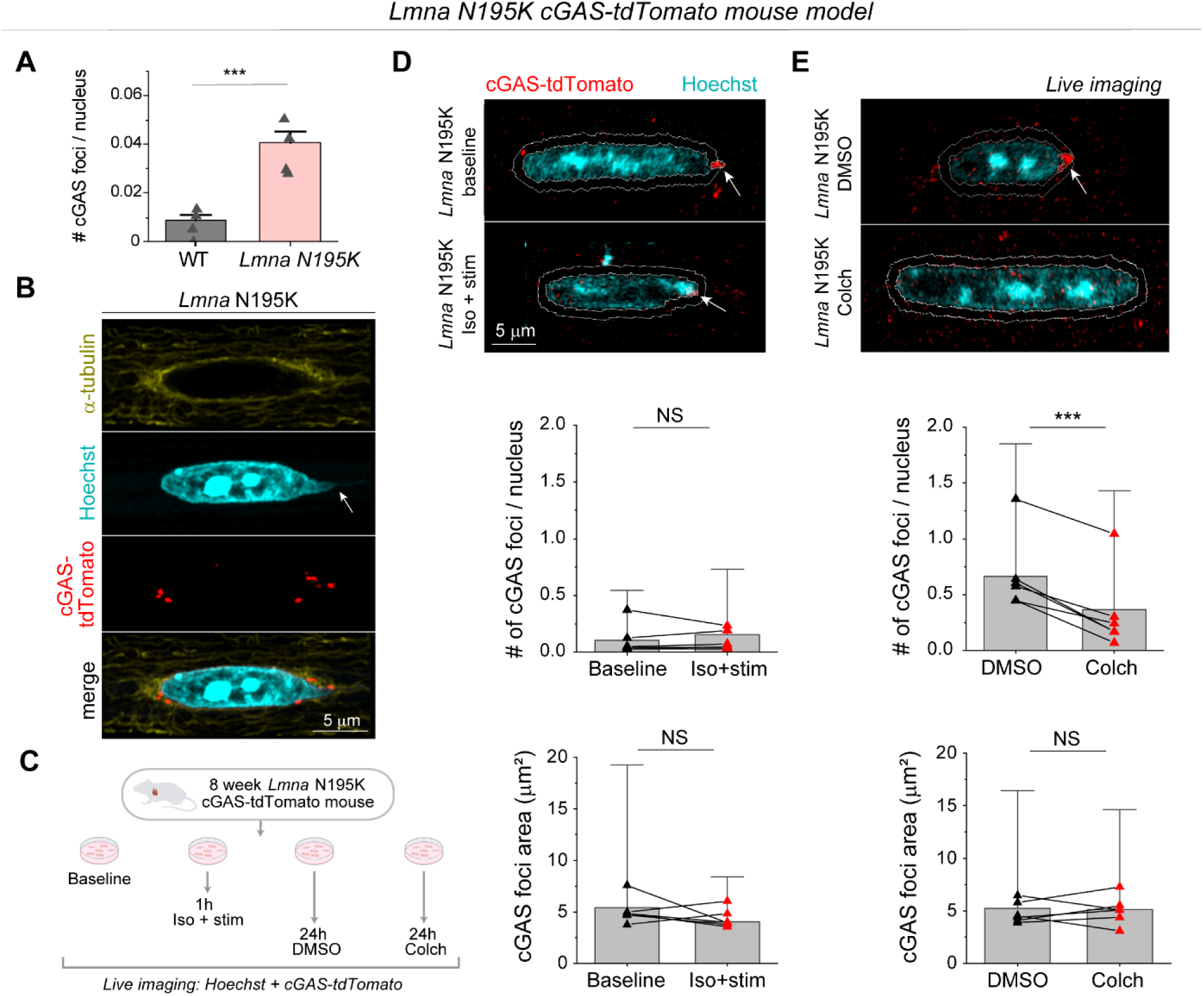
MT disruption protects from nuclear damage in *Lmna* N195K cardiomyocytes. (A) Quantification of number of cGAS foci per nucleus for WT (*N* = 4, *n* = 1898) and *Lmna* N195K (*N* = 4, *n* = 2915) groups. Bar graphs represent the mean ± 1 SD of the pooled nuclei. Superimposed black triangles represent the replicate means. Statistical significance determined by two-tailed t-test. (B) Immunofluorescence mid plane image of the dense MT network (α-tubulin) on the tip of *Lmna* N195K nucleus, associated with a chromatin protrusion (Hoechst, arrow), and cGAS foci (cGAS-tdTomato) indicative of NE ruptures. (C) Experimental design for live imaging of perinuclear cGAS-tdTomato foci in a *Lmna* N195K mouse model, following in-vitro stimulation in the presence of isoproterenol or colchicine treatment. Representative maximum intensity projection images of nuclei (Hoechst) and perinuclear cGAS-tdTomato foci, from (D) freshly isolated (baseline), and contractility induced (Iso + stim) cardiomyocytes, and (E) DMSO or colchicine treated cardiomyocytes. The 1 µm perinuclear rings used for foci identification are indicated with white lines, and the detected cGAS foci are indicated with arrows. Quantification of the number and area of perinuclear cGAS foci, per nucleus (bottom). *N* = 6, *n* = 4869 (baseline), *n* = 2286 (1 h iso + stim), *n* = 4820 (24 h DMSO), *n* = 4263 (24 h colch). Bar graphs represent the mean ± 1 SD of the pooled nuclei. Superimposed black triangles represent the replicate means and connected by lines to their respective treatment conditions. Statistical significance determined by two-tailed *t*-test (*, *p* < 0.05; **, *p* < 0.01; ***, *p* < 0.001).

We first asked whether augmented systolic actomyosin contractility would lead to additional NE ruptures in cardiomyocytes isolated from cGAS-tdTomato *Lmna* N195K mice. We electrically stimulated cardiomyocytes for 1 hour at 0.5 Hz in the presence of isoproterenol to increase contractile workload and compared the number of nuclear cGAS-tdTomato foci to that in non-stimulated cells (Fig. 6C-D). Since the cGAS-tdTomato signal could occasionally be detected in the cytosol (Extended Data Fig. 6B), we restricted our analysis to perinuclear cGAS-tdTomato foci immediately adjacent to the nucleus and within 1 µm of the nuclear border (Fig. 6D-E) to account for small chromatin protrusions difficult to detect based on the Hoechst signal. Augmented systolic workload induced by electrical stimulation and isoproterenol affected neither the number nor size of perinuclear cGAS-tdTomato foci (Fig. 6D), indicating that in this context active sarcomere contractility did not promote NE rupture, possibly due to the limited duration of electrical pacing.

We next tested the alternative hypothesis that MT associated forces drive nuclear damage. We disrupted the MT network in quiescent, isolated cardiomyocytes with 1 µM colchicine for 24 h and compared perinuclear cGAS-tdTomato foci to DMSO treated controls (Fig. 6C). Colchicine treatment induced nuclear elongation, consistent with reduced compressive forces (Extended Data Fig. 6C). MT disruption consistently resulted in a decrease in the number of perinuclear cGAS-tdTomato foci, with no change in their size (Fig. 6E). These data suggest that 24 h of MT disruption prevent the formation of new NE ruptures in *Lmna* N195K cardiomyocytes, without modulating the size of pre-existing ruptures.

We further probed the role of MT forces in a complementary model of *LMNA* deficient, human induced pluripotent stem cell-derived cardiomyocytes (hiPSC-CMs), which exhibit mechanically induced DNA damage ^10,12,13,24,25^. We transfected hiPSC-CMs with *LMNA*-targeted siRNA (siLMNA) or non-targeted siRNA (siNT) for 5 days and disrupted MTs with colchicine on the last day before cell harvesting (see Experimental Design in Extended Data Fig. 7A). Lamin A/C depletion was confirmed by western blot analysis (Extended Data Fig. 7B) and DNA damage was assessed by immunofluorescence for γH2A.X, an early marker for double-stranded DNA breaks ^26^. Extended Data Fig. 7C depicts representative nuclei from hiPSC-CMs, demonstrating increased DNA damage in the *LMNA* depleted nuclei that is significantly reduced upon MT disruption (Extended Data Fig. 7D). Taken together, our findings demonstrate that reducing MT forces in lamin A/C deficient cardiomyocytes protects from nuclear damage.

### MT disruption is sufficient to protect from nuclear damage and preserve cardiac function in mice with cardiac specific *Lmna* depletion

To extend our in vitro observations and test whether MT disruption can protect from nuclear damage and preserve cardiac function upon lamin A/C deficiency in vivo, we used a mouse model with inducible, cardiomyocyte-specific depletion of lamin A/C (*Lmna-*cKO) that we recently generated ^27^. Mice at 10 weeks of age were injected with tamoxifen every other day to induce cardiomyocyte specific *Lmna* depletion^27^. Mice were concomitantly injected with increasing doses of colchicine to disrupt MTs (see schematic for experimental design in Fig. 7A, ascending dose protocol and validation from ^28^). Lamin A/C depletion and MT disruption were confirmed 22 days after tamoxifen induction by western blot analysis (Fig. 7B). At this time point, lamin A/C levels were reduced ∼58%, comparable to recent reports ^15,27,29^, and consistent with the long half-life of lamin proteins. This depletion mimics the reduced lamin A/C levels seen in multiple human *LMNA* cardiomyopathy models ^30,31^. We performed sequential echocardiography analysis at day 0 (before injections), and at day 11, 22, and 29 post injection. At day 22, we observed significantly reduced cardiac function, measured via LV ejection fraction, in the *Lmna-*cKO mice (Fig. 7C), as well as indicators of pathological remodeling (LV dilation and hypertrophy, Extended Data Fig. 8A). Importantly, *Lmna* cKO mice treated with colchicine had fully preserved LV ejection fraction at 22 days (Fig. 7C) and did not show any evidence of chamber dilation or hypertrophy (Extended Data Fig. 8A), demonstrating a significant rescue effect by the MT disruption. Some colchicine-treated mice exhibited reduced cardiac function only at a later point, 29-day post injection, while vehicle-treated *Lmna-*cKO mice did not survive until that time. This longer survival of *Lmna-*cKO mice treated with colchicine is further depicted in the overall extended survival curves (Extended Data Fig. 8B).

**Fig. 7:**
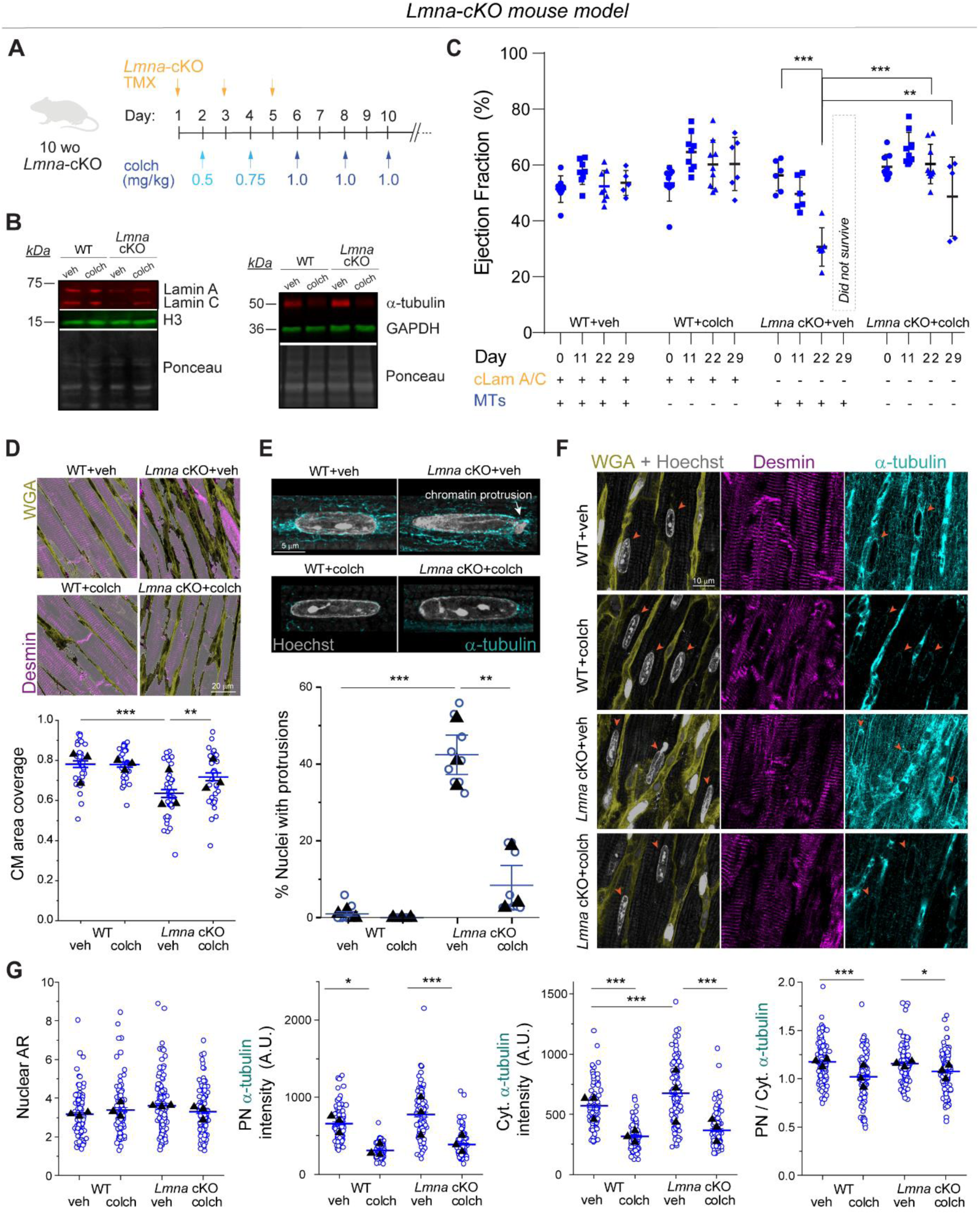
MT disruption preserves cardiac function and protects from nuclear damage in cardiac specific *Lmna* depleted (*Lmna* cKO) mice. (A) Experimental scheme for concurrent MT depolymerization and cardiomyocyte-specific deletion of *Lmna* in mice. Briefly, mice harboring a cardiomyocyte-specific Cre and *Lmna* floxed sites were injected with 3 tamoxifen (TMX) injections at 10-weeks every other day. In between TMX doses, mice were also injected with a ramp-up of colchicine up to 1 mg/kg every other day until sacrifice. (B) Western blot measuring lamin A/C (left) and α-tubulin (right) protein levels 22 days post first TMX injection (C) Sequential % ejection fraction measured by echocardiography in *Lmna* cKO mice with in vivo microtubule disruption. Data shown for pre-injection (day 0), 11, 22, and 29 days post initial injection. Each data point represents 1 mouse (2-3 left-ventricle regions averaged per mouse). WT + veh: *N* = 8 (d29 *N* = 5), WT + colch: *N* = 9 (d29 *N* = 6), *Lmna* cKO + veh: *N* = 6 (*Lmna* cKO + vehicle treated mice do not survive to day 29 post injection), *Lmna* cKO + colch: *N* = 9 (d29 *N* = 5). Error bars represent mean ± 1 SD. Statistical significance determined by two-way ANOVA with Tukey multiple correction. (D) Representative cardiac tissue sections of the indicated groups, stained for Wheat Germ Agglutinin (WGA, yellow) and desmin (magenta). Images are maximum intensity projections of 4 µm thick optical sections. Quantification of CM coverage area, inversely related to fibrosis, from WGA channel. Detected CM areas are highlighted on the images with semi-transparent white. Individual blue circles represent images, black triangles represent replicate means. *N* = 3 mice per group. WT veh: *n* = 35, WT colch: *n* = 37, *Lmna* cKO + veh: *n*=39, *Lmna* cKO + colch: *n*=38 images. Error bars represent mean ± 1 SE. Statistical significance determined by one-way ANOVA with Bonferroni multiple correction. (E) Representative images of cardiomyocyte nuclei (Hoechst, gray) and surrounding MT network (α-tubulin, cyan) of the indicated groups. A chromatin protrusion at the tip of *Lmna* cKO +veh nucleus is indicated with an arrow, with closely encaging MT network. Images are maximum intensity projections of 4 µm thick optical sections. Quantification of the percentage of nuclei with chromatin protrusions in each group by three independent, blinded scorers. Open circles represent the percentage of nuclei with chromatin protrusions quantified by each independent user for each animal. Closed circles correspond to the mean for each animal. Mean ± SE is shown for each group. *N* = 3 mice per group. WT + veh: *n* = 114. WT + colch: *n* = 102. *Lmna* cKO + veh: *n* = 112. *Lmna* cKO + colch: *n* = 113 nuclei. Statistical significance determined by 1-way ANOVA with Bonferroni correction. (F) Representative cardiac tissue sections of the indicated groups, stained for Wheat Germ Agglutinin (WGA, yellow), nuclei (Hoechst, gray), desmin (magenta) and α-tubulin (cyan). Images are maximum intensity projections of 4 µm optical sections. Cardiomyocyte nuclei indicated with arrows. (G) Quantification of cardiomyocyte nuclei aspect ratio (AR), perinuclear (PN), cytoplasmic (Cyt) and PN/Cyt. α-tubulin intensity. *N* = 3 mice per group. WT veh: *n*=113. WT colch: *n*=112. *Lmna* cKO + veh: *n* = 111. *Lmna* cKO + colch: *n* = 112 nuclei. Statistical significance determined by 1-way ANOVA with Bonferroni correction. (*, *p* < 0.05; **, *p* < 0.01; ***, *p* < 0.001).

To further assess cardiac structure, we analyzed cardiac tissue sections at day 29 post injection (for all groups except *Lmna* cKO mice treated with vehicle control, which were analyzed at day 22). We observe decreased cardiomyocyte coverage area, indicative of increased fibrosis, in *Lmna* cKO mice, in agreement with earlier reports ^15,27,29^. Moreover, in vivo colchicine treatment partially restored cardiomyocyte coverage area (Fig. 7D).

To test whether in vivo MT disruption protects cardiomyocyte nuclei from damage, we quantified the percentage of cardiomyocyte nuclei with chromatin protrusions. Similar to the *Lmna* N195K model and an earlier report,^15^ we observed chromatin protrusions specifically at the tips of the *Lmna* depleted cardiomyocyte nuclei, where the perinuclear MT population is enriched (Fig. 7E top). Consistent with a rapid and severe *Lmna*-DCM model, we observed ∼40% of nuclei with chromatin protrusions at day 22 post injection. Notably, in vivo MT disruption robustly protected cardiomyocyte nuclei from chromatin protrusions and damage at day 29 post injection (Fig. 7F), suggestive of potentially even greater effect at comparable day 22. We further characterized nuclear morphology and cytoskeletal organization from cardiac tissue sections and observed no significant changes in nuclear morphology in *Lmna* cKO or colchicine treated mice (Fig. 7F-G, Extended Data Fig. 9A). As expected, in vivo colchicine injections significantly reduced the perinuclear and cytoplasmic MT populations, and specifically reduced enrichment of the perinuclear MT cage (Fig. 7F-G). Colchicine treatment also reduced the density of desmin intermediate filaments, in the WT and *Lmna* cKO mice (Extended Data Fig. 9B). Collectively, these findings are consistent with our observations from the *Lmna* N195K mutant model and demonstrate that disruption of the perinuclear MT cage protects from nuclear damage and is sufficient to preserve cardiac function in *Lmna* cKO mice.

### Computational model of nuclear damage due to resting cytoskeletal forces

To better understand the forces that dictate nuclear morphology and damage, we developed a computational finite element (FE) axisymmetric model for the resting, mature cardiomyocyte (Fig. 8A). In this model, we explicitly consider the nucleus, its surrounding MT cage, and the myofibrils (cytoplasm) as components involved in nuclear deformations. The nucleus is further divided into 1) the nucleoplasm, and 2) the NE and its underlying lamina. Experimental observations suggest a complex prestress field in resting cardiomyocytes that results from cytoskeletal forces that govern nuclear shape and stiffness in these rod-shaped cells ^3,32,33^. To induce this prestress field in our model, we first consider a (imaginary) cylindrical stress-free configuration for the cardiomyocytes where the nucleus is round, and the sarcomere units are not yet assembled (Fig. 8A, Extended Data Fig. 10 and Methods for more details). We further restrict axial displacement (*u_z_* = 0) of the cell ends (Fig. 8A) to mimic the geometric constraints imposed by the myocardial microenvironment ^33^ and restoring stresses from titin springs ^34^. We add an isotropic and homogenous compressive stress field with magnitude *σ_MT_* to the perinuclear MT cage to capture the pushing forces mediated by MT polymerization ^3^ and MT motors (such as kinesin-1) on the nucleus. We assume that in the stress-free configuration, actomyosin fibers are distributed randomly and thus their associated contractility (shown by tensor *ρ_ij_* in our model) is initially isotropic and uniform everywhere in the cytoplasm with magnitude ρ_0_. Note that *ρ_ij_* shows the cardiomyocyte contractility at rest (diastole) ^34^, which differs from its active contractility during systole.

**Fig. 8:**
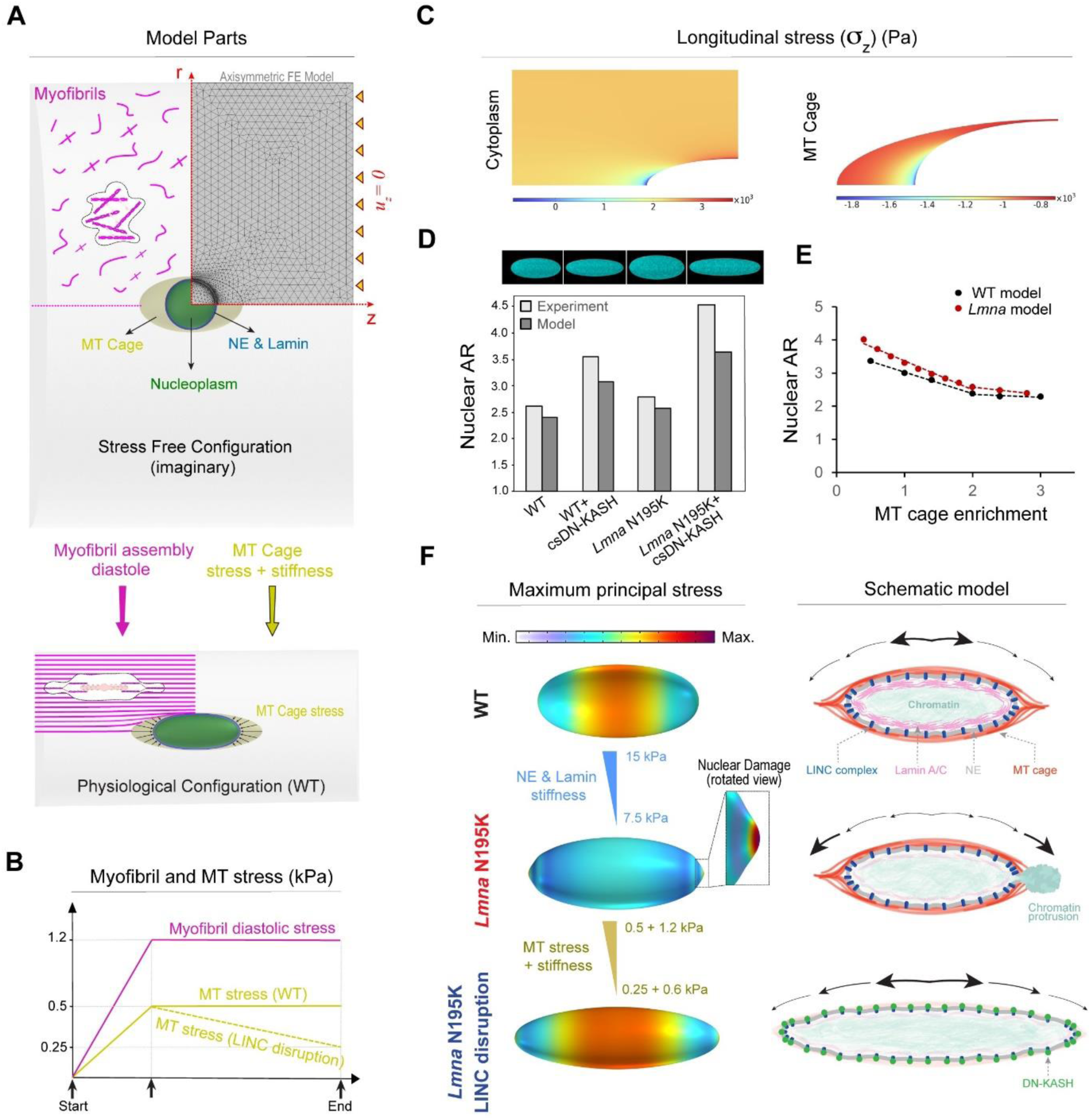
A computational model to explain nuclear damage in resting *LMNA* mutated cardiomyocytes and its rescue through LINC complex disruption. (A) Axisymmetric finite element (FE) model considering an imaginary stress-free configuration (top) for cardiomyocytes consisting of a round nucleus (which is further divided into nucleoplasm, and NE and its underlying lamina), an ellipsoid MT cage, and the surrounding randomly distributed unassembled myofibrils. Diastolic contractility, geometric constraints of the myocardium, restoring forces of titin proteins, and compressive MT forces work in concert to deform this initial configuration to the physiological stressed configuration (bottom) where the nucleus is elongated, and the myofibrils are aligned. (B) Variations of the myofibril diastolic stress and MT compressive stress over the simulation time. (C) Distribution of the longitudinal stress component in the cytoplasm and the MT cage in physiological conditions. (D) Images of simulated nuclei (top) and comparison between the model predictions and the experimental data for nuclear aspect ratio for our four different groups (bottom). (E) Simulation results for variation of the nuclear aspect ratio as a function of MT enrichment for WT and *LMNA* mutated (10 kPa) groups. Closed circles represent simulation data points and dashed lines represent piecewise linear regression fits for each group. (F) Simulations show that beyond a critical value for the MT compressive stresses, a form of instability emerges at the nuclear tips in which the location of maximum principal stress shifts from the middle of the nucleus to the tips (left, top and middle). Continuing the simulation after the instability by reducing MT compressive forces (simulating LINC complex disruption) shows that the nucleus becomes thinner and longer, with the maximum principal stress returning to the middle part (left, bottom). These observations are schematically shown in the right panel.

We next increase ρ_0_and *σ_MT_* from zero (stress-free configuration) to reach the physiologically stressed (WT) configuration (Fig. 8A bottom, Fig. 8B). As a result, the cytoplasm contracts in the radial direction while its length remains constant (Supplementary movie S3), leading to the development of an anisotropic stress field within the cell. This stress field triggers stress-activated signaling pathways which result in polarization of actomyosin fibers ^35,36^ and assembly of myofibrils in the direction of maximum principal stress ^37^. Our simulations show that this principal direction is perfectly aligned with the direction of myofibrils in the cardiomyocytes (Fig. 8A bottom and Extended Data Fig. 10B). Furthermore, our simulations show that myofibril maturation imposes significant lateral compressive forces on the nucleus, leading to its elongation in the axial direction (Fig. 8A bottom). Nuclear elongation is however resisted by elastic compression of surrounding MTs and their pushing forces (*σ_MT_*). Together, these interactions induce the prestress field in the quiescent cardiomyocyte (Extended Data Fig. 10A), even in the absence of cell contractility, although cell contractility might further increase the stress on the nucleus. Our model predicts that maximum and minimum myofibril tensions happen at the short and long tips of the MT cage, respectively, while far from this cage, tension is nearly constant (Fig. 8C). In contrast, maximum and minimum axial MT compression happens at the long and short tips of the nucleus, respectively. The obtained values for diastolic tension of myofibrils (2 − 3 *kPa*) and axial compression of MTs (1 − 1.8 *kPa*) are in good agreement with their corresponding experimental estimations ^32–34^.

We next explored model predictions for nuclear morphology changes due to *LMNA* mutations and LINC complex disruption. To simulate the effect of lamin A/C depletion or mutant lamins, we reduce the stiffness of the NE and lamina ^38,39^. To simulate LINC complex disruption, we decreased both the stress (Fig. 8B) and the stiffness of the MT cage (Fig. 8F), as supported by the loss perinuclear kinesin-1 (Extended Fig. 5D) and MTs (Fig. 5B) upon LINC complex disruption. Our simulations reveal that LINC complex disruption increases the nuclear aspect ratio, which is further exacerbated upon *LMNA* depletion, closely matching our experimental findings (see simulated nuclei images and nuclear aspect ratio comparison in Fig. 8D). This result highlights the significant role of MT enrichment around the nucleus in shaping its morphology. To examine this role in more detail, we then varied MT enrichment in our model by changing the cage stress and stiffness (Extended Data Fig. 10D and methods). Our simulations reproduced the experimentally observed biphasic relationship between nuclear aspect ratio and MT enrichment in the perinuclear region (Fig. 8E).

Our simulations also show that for any specified stiffness of the NE and lamina, a critical value for MT enrichment exists beyond which a form of instability emerges at the nuclear tips (Fig. 8F). Interestingly, the appearance of this instability is concomitant with the shift of the location of maximum principal stress (tension) from the middle of the nucleus to its long tips (Fig. 8F). Recent experimental studies ^40^ indicate that the nuclear lamina can become diluted at the nuclear tips, where the Gaussian curvature is high. Consequently, the combination of high tension and low lamina concentration makes nuclear tips the most likely places to form nuclear lamina gaps, ultimately leading to nuclear damage and rupture ^41,42^. This is consistent with our experimental observation that chromatin protrusions are always observed at the tips. Together, these simulation results suggest that compressive forces of the MT cage are sufficient to drive nuclear damage in quiescent *LMNA*-deficient cardiomyocytes.

Finally, we aimed to simulate rescue of the NE rupture in *LMNA*-deficient cardiomyocytes through LINC complex disruption. After appearance of the instability at the nuclear tips, we simulated LINC complex disruption by decreasing stress and stiffness of the MT cage (Fig. 8B). As a result, the instability disappears, and the nucleus becomes thinner and longer while the location of the maximum principal stress comes back to the middle of the nucleus (Fig. 8F, left). This provides a quantitative explanation for rescue of nuclear damage by LINC complex disruption in *LMNA*-deficient cardiomyocytes.

Overall, our experimental and modeling data suggest that nuclear elongation is regulated by the balance between cross-sectional myofibril compression and the perinuclear MT cage that provides axial compressive forces to resist elongation (Fig. 8F, right). A compromised lamina leads to redistribution of the maximum principal stress to nuclear tips, which contributes to NE rupture and chromatin protrusions. Decoupling the nucleus from MT-based forces via LINC complex disruption or direct MT disruption redistributes the stress away from the tips, thus protecting the nucleus from new ruptures.

## Discussion

This study investigated force transmission into the nucleus during cardiomyocyte contraction and at rest (i.e., non-contracting cells). We describe a novel method to measure active strain transfer from the contracting sarcomeres into the nucleus. Using this and additional approaches, we did not find any evidence to support active sarcomere contractility as a primary driver of nuclear damage in *Lmna*-cardiomyopathy. Instead, we found that nuclear ruptures depend upon the dense cage of perinuclear MTs and associated kinesin motors. Either LINC complex or MT disruption eliminates MT forces and protects from NE ruptures and DNA damage. We further found that in vivo MT disruption protects from nuclear damage, prevents declining cardiac function, and extends survival in mice with cardiomyocyte-specific lamin A/C depletion. Finally, we developed a computational model that demonstrates redistribution of stresses and local vulnerabilities at the long tips of lamin-deficient nuclei, and restoration of stress distribution upon reduction in MT cage stress and stiffness. In sum, our data support a model where local MT compressive forces, independent of actomyosin contractility, induce damage at sites of nuclear fragility in laminopathic cardiomyocytes. Protection from nuclear ruptures and preservation of cardiac function can thus be achieved by disrupting MT-LINC complex interactions.

Our active strain coupling approach provides the first demonstration of sarcomere-nuclear strain coupling in electrically stimulated, mature cardiomyocytes. Earlier studies report nuclear strain in response to passive stretch in non-muscle cells ^1,43^, or spontaneously contracting immature cardiomyocytes ^32,44^ but provided inconsistent insights on cytoskeletal-nucleoskeletal coupling. We report physiologically relevant sarcomere strains of >10%, and high temporal resolution for instantaneous strain coupling with the nucleus, providing amplitude and kinetics for systolic and diastolic phases.

This approach demonstrates that the strain on the nucleus in mature cardiomyocytes is tightly coupled to sarcomere strain during stimulated contraction, but with dampening of nuclear strain amplitude. Surprisingly, this active coupling is only mildly disturbed following perturbation of MTs or the LINC complex. These findings are in agreement with the dispensability of nesprin-1’s actin binding domain on striated muscle structure and function in adult mice ^23^. These results suggests that the central location of cardiomyocyte nuclei facilitates nuclear deformation largely by the surrounding myofibrils that squeeze on the nuclei during contraction, independent of direct physical connection via LINC complexes.

Our study incorporates two complementary approaches to disrupt the LINC complex: 1) acutely in isolated rat cardiomyocytes (48 hrs AdV DN-KASH), and 2) chronically for 5 weeks in mouse hearts in vivo (csDN-KASH). In contrast to the reduced active nuclear strain upon acute LINC complex disruption, chronic in vivo perturbation did not elicit the same effect. This may be due to differential cytoskeletal and nuclear remodeling in response to rapid in vitro vs. chronic in vivo LINC complex disruption. In support of this concept, acute AdV-DN KASH expression led to nuclear expansion in all dimensions and a modest reduction of perinuclear MTs (enrichment ∼1.7-fold, Ext. Data Fig. 5E). Alternatively, chronic csDN-KASH led to more complete reduction of the MT cage (enrichment ∼1.4-fold, Fig. 5B) and marked nuclear elongation. Greater elongation upon csDN-KASH is consistent with the relationship between MT cage enrichment and nuclear aspect ratio measured in Fig. 5C. The lack of expansion in the nuclear short axis upon in vivo csDN-KASH could arise from several factors, including but not limited to, greater nuclear confinement from myofilaments in vivo. Regardless, csDN-KASH nuclei show similarly unaltered strain coupling as with acute MT disruption, consistent with MT disruption driving the csDN-KASH phenotype. Overall, these findings are instructive of nuclear responses to robust and acute LINC complex disruption versus a more chronic and perhaps therapeutically relevant disruption in vivo.

Our strain coupling analysis demonstrates a ∼30% increase in integrated nuclear strain in *Lmna* N195K cardiomyocytes (Fig. 4H). Such increased strain during each contractile cycle may (or may not) be associated with nuclear damage. However, cardiac-specific LINC complex disruption preserves cardiac function and protects *Lmna* N195K nuclei from ruptures independent of reducing active nuclear strain, suggesting increased contractile nuclear strain is not the dominant driver of nuclear damage. In support, increased contractile load did not increase nuclear ruptures (cGAS-tdTomato foci) in laminopathic cardiomyocytes in vitro. However, these results are limited by the short stimulation time (1 hour), and further investigation utilizing chronic alteration in actomyosin contractility is required to conclude on its possible involvement in laminopathy associated NE ruptures in vivo.

Alternatively, our experimental and modeling results demonstrate that MT compressive forces regulate nuclear morphology in the adult cardiomyocyte. During maturation, cardiomyocytes undergo uniaxial elongation, and the centrally located nuclei are squeezed by aligning myofibrils, causing overall nuclear elongation ^33^. The dense perinuclear MT cage is also formed during maturation and provides balancing compressive forces to the nucleus, with a denser concentration at the myofibril void spaces along the longitudinal nuclear tips (Extended Data Fig. 1B). Our analysis demonstrates that nuclear elongation is restricted by the perinuclear MT cage, that when reduced below a critical value (Fig. 5C) leads to nuclear elongation. Consistent with softer and more deformable *Lmna* mutant nuclei, we observe significantly elongated nuclei in LINC complex disrupted (perinuclear MT depleted) *Lmna* N195K cardiomyocytes compared to WT controls. The dramatically elongated nuclei in *Lmna* N195K csDN-KASH mice (where nuclear ruptures are reduced and cardiac function is improved) indicate that elongation of soft nuclei is indeed a common feature of *Lmna* cardiomyopathy, but nuclear elongation itself is not sufficient to drive pathological progression.

When comparing WT and *Lmna* N195K cardiomyocytes with an intact LINC complex (and comparable levels of MT enrichment), we observed no statistically significant difference in nuclear elongation. This differs from earlier reports on elongated nuclei in alternative laminopathy mouse models ^7,8^. This discrepancy can be explained by differences in cytoskeletal remodeling or disease severity between the models but might also be partly influenced by imaging resolution and selection criteria. Here we employed high resolution imaging that allows identification of nuclei with chromatin protrusions, which we excluded from nuclear shape analysis (20% in *Lmna* N195K group). Indeed, we observe mild (11%) but statistically significant elongation of *Lmna* N195K nuclei on a larger data set with lower resolution imaging and no excluded nuclei (Extended Data Fig. 2B). Further, our observation of chromatin protrusions and nuclear ruptures at the tips of the nuclei, without significant nuclear elongation, agree with recent reports from *Lmna*-depleted mature cardiomyocytes correlating nuclear ruptures with phenotype severity ^15,29^.

We demonstrate the loss of perinuclear MTs and interacting kinesin-1 motors upon cardiac-specific LINC complex disruption in a cardiac laminopathy model, concomitant with a reduction in NE damage. We previously demonstrated a protective role of kinesin-1 depletion in *Lmna-*deficient skeletal myotubes, where MT motors are required for nuclear migration during myotube maturation ^10^. In maturing skeletal muscle myotubes, nuclear damage is driven by cytoskeletal forces associated with kinesin-1 dependent nuclear movement. However, the cardiomyocyte nuclei remain centrally located in maturation, and the damage in *Lmna-* compromised nuclei appears to be driven by compressive forces from the perinuclear MT cage. Despite this mechanistic distinction, convergent strategies that disrupt kinesin’s interaction with the nucleus warrant further examination for striated muscle laminopathies.

Computational modeling provides additional insights into the subcellular forces underlying nuclear damage and rescue. The model incorporates tensile forces from maturing myofibrils, stiffness of the nuclear lamina, and combined stress and stiffness from the MT cage and associated motors. Simulations demonstrate the redistribution of overall nuclear stress to the long nuclear tips in laminopathic nuclei with intact MT forces, localizing a site of nuclear vulnerability. The localized stress at the nuclear tip is restored to the center of the nucleus when both MT stiffness and stress are reduced, concomitant with nuclear elongation. This provides an important quantitative framework to support the sufficiency of resting, perinuclear MT forces for nuclear damage upon lamina compromise.

We incorporated an in vivo mouse model with inducible, cardiomyocyte-specific *Lmna* depletion to test the potential of MT disruption in protecting compromised nuclei and preventing deterioration of cardiac function. This rapid and aggressive *Lmna* cardiomyopathy model presents significant cardiac dilation and reduced function within 3 weeks, with reduction in lamin A/C levels comparable to reports from *LMNA* cardiomyopathy patients ^27^. Our cellular analysis identified chromatin protrusions in ∼40% of cardiomyocyte nuclei, in agreement with recent studies utilizing this approach that reported nuclear abnormalities and ruptures preceding reduced cardiac function ^15,29^. Accordingly, our experimental design induced concomitant *Lmna* depletion and MT depolymerization. In support of our findings in *Lmna* N195K mice, we identify chromatin protrusions at the tips of the elongated cardiomyocyte nuclei, closely encaged by the perinuclear MTs. These perinuclear MTs can be significantly reduced by in vivo colchicine administration, thus protecting the nuclei from damage, improving cardiac function and remodeling, and extending lifespan in *Lmna* depleted mice. Despite the reduced perinuclear MT cage, we did not observe significant nuclear elongation in the WT or *Lmna*-depleted nuclei treated with colchicine, which might be attributed to distinct force profiles acting on the nucleus in vivo vs. ex vivo, and/or distinct cytoskeletal remodeling in the faster and more severe *Lmna*-depletion model, compared to longer and milder *Lmna* N195K model with LINC complex disruption. The latter notion is partially supported by colchicine-induced reduction in cytoplasmic desmin (Extended Data Fig. 9B). While these results demonstrate the therapeutic potential of targeting MTs in cardiomyocytes in patients with *LMNA* cardiomyopathy, we have not directly tested the ability of MT disruption to rescue cardiac function in already developed cardiomyopathy. Moreover, the tolerable colchicine doses in mice are two orders of magnitude higher than the tolerable dose in humans. Therefore, transferring this potential safely to the clinic will require perturbing specific MT interactions with the nucleus, with doses that will not interfere with other crucial functions of MTs in the cell.

Currently, a wide range of experimental models is available to study cardiac pathogenesis in laminopathies, including *Lmna*-depletion mouse models, *Lmna*-mutant knock-ins representing several identified patient mutations, and human iPSC-derived cardiomyocyte (iPSC-CM) models ^45,46^. In the present study, we span the above models and demonstrate the importance of studying mature cardiomyocytes with their unique cytoskeletal and nucleoskeletal organization, to identify drivers of nuclear damage in the pathogenesis of laminopathy-associated cardiomyopathies. Although a powerful tool, the human iPSC-CM model represents immature cardiomyocytes. Despite evidence for *LMNA* induced DNA damage, they lack nuclear ruptures likely due to an immature cytoskeleton and nucleoskeleton, and as a result exhibit rounder nuclear morphology with altered stress distribution on the nucleus, which limits the utility of this model to study *LMNA* associated nuclear damage ^12,25,47,48^. Within the mature mouse models, systemic or cardiomyocyte specific *Lmna* depletion is the most aggressive and rapidly progressing DCM model. Mouse models with specific mutations such as the *Lmna* N195K and *Lmna* H222P present variable severity phenotypes, but ultimately all lead to progressive DCM with contractile disfunction and fibrosis. Nuclear structure abnormalities are a common theme in all mature *Lmna* cardiomyocytes, but the wide spectrum of nuclear abnormalities, and accompanied cytoskeletal remodeling, complicate mechanistic insights on disease pathogenesis. Elongated nuclei morphology is a feature of *Lmna* H222P model ^8,49^, which also displays decreased acetylated MT population ^50,51^. In comparison, here we report chromatin protrusions in *Lmna* N195K nuclei, which are elongated (and protected from protrusions) upon elimination of perinuclear MT cage. Moreover, we observe even higher chromatin protrusion rate in the *Lmna* depletion model, in agreement with a recent report^15^. Importantly, in vivo disruption of MTs is sufficient to protect from the severe nuclear damage with lamin A/C depletion supporting a unifying mechanism for MT forces in driving nuclear damage in *Lmna* cardiomyopathy. Since nuclear stability is only mildly impaired in *Lmna* H222P muscle cells, unlike in *Lmna* N195K and *Lmna* deletion models ^10^, it will be imperative to test whether LINC complex or MT disruption also improves cardiac function or survival in the Lmna H222P animals. Additionally, further characterization of the MT network in the *Lmna* H222P model will be beneficial to identify potential adaptive cytoskeleton remodeling mechanisms that might decrease the rate of DCM progression in this model.

### Limitations of the current study

This study focused on the role of cytoskeletal forces as a pathological driver in *Lmna* cardiomyopathy. While the role of the nuclear lamina in chromatin organization and gene expression was outside the scope of this study, it is also hypothesized to participate in the molecular pathogenesis of laminopathies ^14,52,53^. The dual role of lamin A/C in providing nuclear integrity but also organizing chromatin at the nuclear periphery, which in turn affects nuclear mechanical stability, points to potentially integrated effects of structural stability and chromatin organization in driving *LMNA* pathology ^54^. While our study implicates resting MT forces, but not actomyosin contractility in laminopathy nuclear damage, further studies with chronically increased contractility are required to rule out potential involvement in contractility-driven damage to the nucleus, particularly in an in vivo setting. Further, the AdV empty construct, that was used as a control for acute AdV DN-KASH transductions, does not rule out other potential functions of KASH overexpression at the NE. Our cardiac function analysis in *Lmna* N195K csDN-KASH mice demonstrates preserved cardiac function at 10-and 12-week time points. Since these mice eventually perish at 17 weeks, we cannot rule out non-cardiac causes as contributors to mortality. Similarly, we observe extended survival for *Lmna* cKO mice treated with colchicine; however, colchicine treatment by itself is toxic and contributes over time to the mortality of the animals, confounding the analysis. Due to the long-term toxicity of colchicine treatment, we were unable to test its efficacy in improving cardiac function or survival in the Lmna N195K model, which would require extended treatment periods. Given these limitations of colchicine treatment, we primarily consider these studies as proof-of-principal studies to demonstrate the crucial role of the perinuclear MT cage in the pathogenesis of LMNA cardiomyopathies and as a potential therapeutic target. For clinical applications, though, we postulate that more refined treatment approaches would be needed instead of complete MT disruption.

In summary, our study introduces a novel method to investigate active sarcomere-nuclear strain coupling in the primary beating cardiomyocyte and implicates compressive MT forces in nuclear damage in laminopathy. We conclude that targeting specific interactions between MTs and the LINC complex should be pursued as a potential cardioprotective strategy in *LMNA* cardiomyopathy.

## Acknowledgements

Funding for this work was provided by the National Institute of Health (NIH) R01s-HL133080 and HL149891 to B.P., R01 HL082792, R01 AR084664, and R35 GM153257 to J.L., and R35 HL16663 to R.J., the Foundation Leducq Research Grant no. 20CVD01 to B.P. and J.L., the Foundation Leducq Research Grant no. 24CVD03 to J.L., and by the Center for Engineering Mechanobiology to R.J., B.P., and V.S. through a grant from the National Science Foundation’s Science and Technology program: 15-48571. Funding was also provided by the National Science Foundation (CBET 1715606 and URoL-2022048 to J.L.), and the Volkswagen Foundation (A130142 to J.L.), as well as a Fleming Research Fellowship to J.H. We thank the Bridge Position Program, Weizmann Institute of Science for support to D.A.P. We thank R. Rotkopf from the Life Sciences Core Facility, Weizmann Institute, for advice on statistical analysis. We thank Justin Bi for his help as a blinded scorer in the chromatin protrusion analysis. The Lammerding lab thanks the Mills family for their generous support and the Cornell CARE staff, Ern Hwei Hannah Fong and Alexander Sopilniak-Mints for their invaluable contribution to the maintenance of our animal colony.

## Methods

### Animals

Animal care and use procedures were performed in accordance with the standards set forth by the University of Pennsylvania Institutional Animal Care and Use Committee and the Guide for the Care and Use of Laboratory Animals published by the US National Institutes of Health. Protocols were approved by the University of Pennsylvania Institutional Animal Care and Use Committee. All animals provided by the Lammerding Lab at Cornell were bred and maintained according to relevant guidelines and ethical regulations approved by the Cornell University Institutional Animal Care and Use Committee, protocol 2011-0099. Both rats and mice were housed in a facility with 12-h light/dark cycles and provided ad libitum access to water and chow. Temperature and humidity were checked daily to ensure that these parameters stay within appropriate ranges (20–26 C°, 30–70%, respectively). *α*MHC Cre, KASH (csDN-KASH) and *Lmna*^N195K/N195K^ mice have been described previously ^9,19^ and were back-crossed at least seven generations into a C57BL/6 line ^10^. To generate *α*MHC Cre^+/-^ KASH^+/-^ *Lmna*^N195K/N195K^ mice (*Lmna* N195K csDN-KASH), male *α*MHC Cre^+/-^ *Lmna*^N195K/+^ and female KASH^+/-^ *Lmna*^N195K/+^ mice were crossed to create experimental and control littermate mice. All mice were maintained in a uniform C57BL/6 background. *Lmna* mutant mice were provided with gel diet supplement (Nutri-Gel Diet, BioServe) to improve hydration and overall health and quality of life. To induce cardiomyocyte specific KASH-mediated LINC complex disruption, 30mg/kg tamoxifen (Cayman chemicals cat#13258) suspended in sunflower oil (Sigma cat# 88921) was injected intraperitoneally daily for five consecutive days, starting at approximately 3 weeks of age, followed by a 1 week wash out. Controls included *α*MHC Cre^+/-^ KASH^+/-^ *Lmna*^N195K/N195K^ mice treated with vehicle and *α*MHC Cre^+/-^ KASH^+/-^ *Lmna*^+/+^ mice littermates.

To generate *Lmna*^N195K/N195K^ mice expressing cGAS-tdTomato, the previously described cGAS/MB21D1-tdTom transgenic mouse ^10^ was crossed into the *Lmna N195K* background to generate 3×FLAG-cGAS^E225A/D227A^-tdTomato positive *Lmna*^N195K/N195K^ mice within two generations. All mice were maintained in a uniform C57BL/6 background. Single cardiomyocyte data was collected at stated timepoints. Controls included *Lmna^+^*^/+^ mice expressing the cGAS construct.

To generate the inducible, cardiac specific *Lmna* deletion mouse model (*Lmna-*cKO), αMHC-MerCreMer (The Jackson Laboratory (JAX), strain 005657) and *Lmna* floxed mice (JAX strain 026284) were crossed, as previously described ^27^, to obtain *α*MHC Cre^+/-^ *Lmna*^fl/fl^ mice. αMHC-Cre^+/-^ *Lmna*^+/+^ littermates served as controls (*Lmna*-cWT). All mice were maintained in a uniform C57BL/6 background. To induce lamin A/C depletion in cardiomyocytes, tamoxifen was dissolved in sunflower oil (Sigma cat# 88921) to a concentration of 30 mg/kg and injected intraperitoneally (IP) at 10 weeks of age every other day for three total injections (days 1,3 and 5). Vehicle-only injections served as additional controls. To disrupt MT in-vivo mice were also injected in between tamoxifen doses with increasing doses of colchicine (days 2,4,6,8,10 sacrifice; see concentration in Fig. 6F). PBS of equal volume was IP injected for controls. Lamin A/C depletion was validated by western blot on cardiac tissue obtained from mice at 22 days post initial tamoxifen injection, using the anti-lamin A/C (sc-376248, 1:1000) and the anti-H3 (CST #44995, 1:5000) as loading control. MT disruption in the heart was validated by western blot using anti α-tubulin (ab7291, 1:3000) and GAPDH as loading control (CST #2118S, 1:5000).

### Echocardiography

Mice were anesthetized with 2% isoflurane and placed on a stereotactic heated scanning base (37°C) attached to an electrocardiographic monitor. Left ventricular structure and function were determined with a Vevo 2100 imaging system (VisualSonics) equipped with a MS550D transducer (22–55 MHz). Echocardiographic parameters were measured for at least five cardiac cycles using AM mode images. Analysis was performed using the AutoLV Analysis Software (VisualSonics) and conducted by observers blinded to the mouse genotype and/or treatment groups.

### Adult rat and mouse cardiomyocyte isolation and culture

Primary adult ventricular myocytes were isolated from 8-to 12-week-old Sprague Dawley rats, or 8-9 week-old mice using Langendorff retrograde aortic perfusion with an enzymatic solution as previously described ^21^. Briefly, the heart was removed from an anesthetized rodent under isoflurane and retrograde-perfused on a Langendorff apparatus with a collagenase solution. The digested heart was then minced and triturated with glass pipettes to free individual cardiomyocytes. The resulting supernatant was separated and centrifuged at 300 rpm to isolate cardiomyocytes. These cardiomyocytes were then resuspended in cardiomyocyte media (Medium 199 (Thermo Fisher) supplemented with 1x insulin-transferrinselenium-X (Gibco), 1 μg/μL primocin (InvivoGen), and 20 mM HEPES, pH = 7.4 (UPenn Cell Center)) at low density, cultured at 37 C° and 5% CO2 with the addition of 25 μmol/L of cytochalasin D in the media.

### Active sarcomere-nuclear strain coupling

Active strain coupling was quantified by back-to-back measurement of sarcomere length contractility and nuclear deformation during electrically stimulated contractions in adult cardiomyocytes (no blebbistatin is added to allow for actin-myosin contractility). Cardiomyocytes in culture media were loaded with 4µM Hoechst and transferred to a custom-fabricated cell chamber (IonOptix) mounted on an LSM Zeiss 880 inverted confocal microscope with 63×oil 1.4 numerical aperture objective. Experiments were conducted at room temperature, and field stimulation was provided at 1 Hz with a cell stimulator (MyoPacer, IonOptix). Rod shaped cells with stable contractions were selected, and baseline sarcomere length >1.7µm, and sarcomere length strain > 10% were used as inclusion criteria. For each cell, sarcomere length contractility (SL) was first measured with a transmitted light camera (IonOptix MyoCam-S) and real time optical Fourier transform analysis (IonWizard, IonOptix). For each cell, 5 steady state and consistent SL traces were recorded in a region of interest close to the nucleus, either above or below the nucleus. The microscope was then immediately switched to fast mode Airyscan confocal and Hoechst fluorescence was imaged with 405nm laser (lowest laser power), at 11 msec/frame (91 Hz), for 5 additional steady state contractions. Initial image analysis was performed using ZEN black software for Airyscan processing, which involves signal integration from the 32 separate sub-resolution detectors in the Airyscan detector and subsequent deconvolution of this integrated signal. Stimulation times were recorded with the sarcomere contractility and nuclear imaging files for offline alignment of the traces. An average sarcomere contractility trace was generated for each cell (IonWizard, IonOptix) and exported to Matlab (Mathworks R2022b) for further alignment with nuclear deformation traces. Nuclear image analysis was performed with Arivis V4D 4.0-4.1, by a pipeline to auto-segment (Otsu) the nucleus object in each frame and export nuclear morphology parameters. Nuclear length and width traces were imported to Matlab, averaged for each cell, and aligned with the corresponding average sarcomere contractility trace (after interpolation to 91 Hz to match nuclear time trace). Sarcomere and nuclear strains were calculated by dividing the corresponding instantaneous strains by the baseline lengths prior to stimulation. For each experimental group 15-20 cardiomyocytes were recorded, from 3-4 biological replicates.

### hiPSC-derived cardiomyocytes

Human induced pluripotent stem cell-derived cardiomyocytes (hiPS-CMs) were purchased from Ncardia (Nc-C-BRCM). Cells were maintained in RPMI supplemented with 10% FBS (Gibco, 16000044), 1% Penicillin-Streptomycin (Gibco, 15140122), and 2% B-27 (Gibco, 17504044). After 4 days in maintenance media, 150,000 cells were plated onto glass coverslips (for immunofluorescence) or into 12-well culture dishes coated with 10 μg/ml fibronectin (Sigma-Aldrich, F1141) diluted in DPBS with Mg^2+^ and Ca^2+^. Cells were then allowed to recover for 4 days in maintenance media. On day 0, cells were treated with siRNAs, against either a non-targeting control (siNT) or lamin A/C (siLMNA) at a concentration of 5nM using Lipofectamine RNAiMax transfection reagent diluted in Opti-MEM. Cells were incubated with siRNAs for 5 days total, and media was changed on days two and four. On day 4, cells were also treated with either DMSO as a control, or colchicine at a concentration of 1µM. 24 hours after DMSO or colchicine treatment, cells were harvested for downstream analysis.

### Immunofluorescence of isolated cardiomyocytes

Primary rat and mouse cardiomyocytes were fixed in 4% PFA (Electron Microscopy Sciences) for 10 min, washed three times with PBS, and permeabilized in 0.1% TritonX-100 for 10 min at room temperature. After washing twice with PBS, cells were placed in blocking buffer (1:1 Seablock (Abcam) and 0.1% TritonX-100 (Bio-Rad) in PBS for at least 1 h at room temperature, then labeled with primary antibodies (see below) for 24–72 h at 4 °C. Cells were then washed three times in PBS, then labeled with secondary antibodies in PBS at room temperature for 2–4 h. Hoechst was added for the last 10 min of secondary incubation and cells were washed twice with PBS. Stained cells were mounted on #1.5 coverslips in Prolong Diamond Antifade Mountant (Thermo Fisher) for imaging. Slides were left to cure in the dark for at least 24 h prior to imaging. hiPS-CMs coverslips were fixed in 4% PFA for 10 minutes at room temperature, then rinsed in DPBS for 5 minutes three times. Cells were permeabilized with 0.5% TritonX-100 for 10 minutes, blocked in 1% BSA in PBS-T (8mM Na_2_HPO_4_, 150mM NaCl, 2mM KH_2_PO_4_, 3mM KCl, 0.05% Tween 20, pH 7.4) and incubated with primary and secondary antibodies diluted in PBS-T + 1% BSA for 1 hour each at room temperature. Samples were counterstained with DAPI solution (Sigma, D9542) for 10 minutes at room temperature, then rinsed with PBS and stored/imaged in SlowFade Gold Antifade Mountant (Invitrogen, S36936).

### Mouse tissues processing and staining

Immediately following tissue collection, hearts were washed with 1X PBS and fixed in 4% paraformaldehyde (diluted in 1X PBS) at 4°C overnight. For immunofluorescence analysis, samples were washed with PBS and flash frozen in Tissue-Tek® O.C.T. Compound (Sakura, #4583) in an ethanol bath, followed by storage at –70°C. Frozen tissue blocks were cryosectioned using an Epredia™ Microm HM525 NX Cryostat (SN: S20020277) to a thickness of 5 μm, mounted on 25 × 75 × 1.0 mm Superfrost Plus Microscope slides (Fisherbrand® #12-55015) and left to air-dry for 1 hour, followed by blocking and permeabilization with a solution of 3% BSA, 5% filtered horse serum in PBS-T (0.05% Triton-X 100 and 0.03% Tween (Sigma) in 1X PBS) for 1 hour at room temperature.

For picrosirius red staining, whole ventricles were cryosectioned at 5 μm intervals and left to air dry for at least 1 hour at room temperature and stored at-20 C. Upon processing, slides were brought to room temperature and put in Bouin’s fixative (Electron Micrscopy Sciences #15990-10) set in a 56 C water bath for 15 minutes then washed 2x with milliq water. Slides were then submerged in Sirius red (Sigma #365548-5G, Direct Red 80 in 1.3% picric acid, Sigma #P6744-1GA) for 2 hours gently rocking at room temperature. Slides were washed 2x in 0.5% acetic acid then dehydrated using 95% then 100% ethanol. Finally, slides were equilibrated in xylene substitute and mounted in a xylene-based media (Leica #3801731) and imaged using ScanScope Cs2 from Aperio and analyzed using the color deconvolution macro in Aperio’s analysis software package.

### Immunofluorescence of cardiac tissue sections

5-µm-thick sections were placed on glass slides and processed for immunofluorescence as described in ^55^ with modifications. The paraffin wax was melted and deparaffinization was achieved by 2×20 min washes in xylene (StatLab, 8400-1). Samples were rehydrated through sequential 2 min washes in decreasing concentrations of ethanol (100%, 95%, 75%, 50%) followed by ddH2O. Then, glass slides were placed in 1x reveal decloaker solution (Biocare Medical, RV1000M) and heat antigen retrieval was performed for 15 min at high pressure using a pressure cooker (Instant Pot Pro 10-in-1 pressure cooker, 8 Quart, Amazon.com). After cooling, residual reveal decloaker solution was washed off in ddH2O and a hydrophobic ring was hand drawn surrounding the sections using a Super PAP pen (Electron Microscopy Sciences, 71312). Sections were washed in 1X PBS and permeabilized with 1X PBS + 0.25% Triton X-100 for 10 min. After permeabilization, sections were thoroughly washed in 1X PBS. An additional wash in 0.1% PBST (0.1% Tween 20 in 1X PBS) preceded blocking for 1 h at RT in Blocking Solution (3% BSA (Sigma Aldrich, A7906) in 0.1% PBST).

Incubation in primary antibodies (Mouse monoclonal to α-tubulin, Sigma Aldrich T5168, clone B-5-1-2, 1:50; Goat polyclonal to desmin, R&D Systems AF3844, 1:250) was performed in Blocking Solution for 48 h, at RT, in a home-made humidified chamber. Sections were rinsed in 1X PBS, and washed 2×15 min with 0.1% PBST. Then, incubation in secondary antibodies (donkey-anti-goat IgG Alexa Fluor 647, Invitrogen A21447, 1:100; donkey-anti-mouse IgG Alexa Fluor 568, Invitrogen A10037, 1:100) was performed in 0.1% PBST for 48 h, at RT, protected from the light in the humidified chamber. Alexa Fluor 488 conjugated WGA (Invitrogen, W11261) was added during the secondary antibody incubation (1:500, final concentration 25 μg/mL). The sections were rinsed in 1X PBS and stained for 20 min with Hoechst 33342, Trihydrochloride, Trihydrate (Invitrogen, H3570, final concentration 10 μg/mL in 1X PBS). Sections were thoroughly washed in 1X PBS before mounting in Prolong Diamond Antifade Mountant (Invitrogen, P36961).

### Antibodies, labels, and pharmaceuticals

Hoechst 33342 (Invitrogen, H3570); SPY650-DNA, SPY555-Tubulin, and SiR-actin (0.1µL/mL, Spirochrome); Anti-alpha Tubulin mouse monoclonal antibody, clone DM1A (1:500 abcam ab264493); Anti-KIF5B rabbit monoclonal antibody clone EPR10276(B) (1:500 abcam ab167429); Anti-Nesprin1 rabbit monoclonal antibody clone EPR14196 (1:250 abcam ab192234); Anti-Nesprin-2 (1:300, kind gift from Hodzic lab); Anti-Lamin A/C (1:1000, Santa Cruz sc376248; Anti-Lamin A/C mouse monoclonal antibody clone 4C11 (1:500, CST #4777); phospho-H2A.X (1:1000 05-636, Millipore Sigma); Anti-Desmin goat polyclonal antibody (1:250 R&D Systems AF3844), Anti-Desmin (1:1000, PA-1151113); Anti-MHY7 (1:250, DSHB BA-D5); Goat anti-rabbit IgG AF 647 (1:1000, Life Technologies, A27040); Goat anti-mouse IgG AF488 (1:1000, Life Technologies, A11001); Colchicine (1 μM in DMSO, Sigma-Aldrich); Isoproterenol (1 μM in DMSO); Blebbistatin (10 μM in DMSO, Cayman Chemical).

### Image acquisition

#### Live 3D imaging – resting primary cardiomyocytes

For live 3D super-resolution imaging adult rat cardiomyocytes in culture media were loaded overnight with 0.1µL/mL SPY650-DNA (Spirochrome) to label nuclei, and 10µM Blebbistatin immediately before imaging, to prevent motion artifacts. Airyscan SR (super resolution) z-stacks were acquired with Zeiss 880 Airyscan confocal microscope with 63×oil 1.4 numerical aperture objective, and 640nm laser line in a glass bottom dish. Raw images were processed with a joint deconvolution plugin (ZEN 3.5 blue).

#### Live cGAS-tdTomato foci – resting primary cardiomyocytes

Live isolated cGAS-tdTomato cardiomyocytes were adhered to glass bottom dishes using MyoTak (IonOptix) and loaded with 4µM Hoechst and 10µM Blebbistatin immediately before imaging. Tile scan images were acquired at room temperature with Zeiss 980 Airyscan confocal microscope equipped with Plan-Apochromat ×20 air 0.8 numerical aperture objective. Two channels (tdTomato and Hoechst), 1 μm z-stacks tile scans were acquired, stitched and Airyscan processed using ZEN black software. Nuclei from dead or severely deformed cells, or with motion or stitching artifacts were excluded from analysis.

#### hiPSC-CM imaging

Z-stacks (10 µm range with voxel size of 0.035 x 0.035 x 0.5 µm) of individual nuclei were acquired on a LSM Zeiss 980 Airyscan 2 confocal microscope. A Plan-Apochromat 63×oil 1.4 numerical aperture objective was used. Tile imaging was performed under 4x zoom and 2x bidirectional averaging. The SR-4Y Multiplexing (MP) acquisition mode was used for faster parallel pixel readout. Samples were excited with the 405 nm, 488 nm and 639 nm laser lines. For nuclei selection, the Hoechst channel was used to preview images and 33.7 x 33.7 µm regions of interest (ROI) were defined to frame individual nuclei. To ensure examination of properly individualized hiPSC-derived cardiomyocytes, ROIs containing more than one nucleus were not selected for imaging. Images were Airyscan processed using the ZEN black software.

#### Mouse tissue sections

Imaging was performed on ≥2 myocardial areas from each of at least 3 individual sections of every animal. Areas with clear striations corresponding to longitudinally sectioned cardiomyocytes were identified through the eyepiece. Z-stacks (6x 1 μm slices, 6 μm range) were acquired on a LSM Zeiss 980 Airyscan 2 confocal microscope using a Plan-Apochromat 63xoil 1.4 numerical aperture objective. Imaging was performed under 1.7x zoom and 2x bidirectional averaging, with 0.035 x 0.035 μm pixel size. Images were acquired in Super Resolution acquisition mode at 2.0x Nyquist sampling. The 639 nm, 561 nm, 488 nm and 405 nm laser lines were used for imaging of desmin, α-tubulin, WGA and Hoechst, respectively. Images were Airyscan processed using the ZEN black software.

### Image analysis

Raw images were Airyscan processed with ZEN Black software and imported to Arivis V4D 4.0-4.1 for further analysis. Dedicated V4D analysis pipeline was generated to auto segment (otsu) the nuclei from 3D z-stacks or 2D maximum intensity projection (MIP) images, and export nuclear morphology parameters, such as nuclear volume/area, length, width, and aspect ratio.

#### Chromatin protrusions

To determine the percentage of nuclei with chromatin protrusions for each group in primary adult mouse cardiomyocytes or mouse tissue sections, three individuals were recruited to do a blind analysis. In consistency with a recent report, nuclear were identified as bearers of chromatin protrusions when having both 1) an external outline deviating from the resting ovoid shape and 2) strong DNA condensation at the nuclear poles protruding from the expected nuclear ovoid shape, leading to nuclear deformation. In the case of the mouse tissue sections, only nuclei with a discernible outline within the z-stack were included in the analysis. A stack of 819 (in the case of primary mouse cardiomyocytes) or 441 (for mice tissue sections) maximum intensity images of Hoechst-stained nuclei from all animals was given to each individual scorer to categorize as nuclei with or without a chromatin protrusion according to the above-mentioned criteria. The average chromatin protrusions from each animal were defined from the percentage of chromatin protrusions scored by each blinded user. The average chromatin protrusions for each group were in turn derived from the average scores of individual animals of the corresponding group.

#### Perinuclear enrichment in isolated CMs

Perinuclear MT or Kinesin-1 enrichment was calculated on MIP images from 3×1µm z-stacks slices (total 3µm) around the midplane of the nucleus. A perinuclear ring object was created from 0.5µm dilation of the nuclear object and subtraction of the nucleus. Nuclear pole enrichment was calculated from manually traced 2 µm wide objects for each of the long and short nuclear poles.

#### cGAS-tdTomato foci

cGAS foci were calculated from tile MIP images with auto segmentation (Otsu) of the nuclei from the Hoechst channel, and generation of 1 µm perimeter perinuclear ring. Fixed intensity (>600AU, or >200AU for the WT vs *Lmna* N195K cGAS-tdTomato models) and size (>0.8µm^2^) thresholds for cGAS-tdTomato signal were used to segment the cGAS foci in the vicinity of the perinuclear ring. The segmented cGAS foci were used to quantify the percentage of cGAS positive myocytes. To prevent contamination by cytosolic cGAS intensity, only foci touching the Hoechst signal and originating within the 1 µm diameter perinuclear rings were included as readouts for nuclear ruptures.

#### γH2A.X foci

hiPSC-CMs images were analyzed with custom Arivis V4D 4.1 pipeline to segment the Hoechst channel using the Otsu method to outline the nuclei in 3D z-stacks. γH2A.X foci were segmented using fixed intensity (>2000 A.U.) and volume (0.02-1000 µm^3^) thresholds. Nuclear volume and mean nuclear intensity of γH2A.X signal were calculated using the nuclear segmentation. Puncta volume, integrated volume of all intranuclear puncta, and number of puncta were calculated from the segmented γH2A.X images. The data were imported to Matlab (Mathworks R2022b) for pooling, and subsequent graphing and analysis was performed using Origin 2019 (OriginLab Corporation). The integrated volume of all γH2A.X foci within a nucleus was divided by the corresponding nuclear volume to calculate the fraction of the nuclear volume occupied by γH2A.X foci, or foci fraction of volume coverage. The integrated volume of all γH2A.X foci within a nucleus was divided by the number of foci in that nucleus to calculate the mean γH2A.X foci volume. Data points with a value greater than 3 standard deviations (SD) above the mean of the γH2A.X mean foci volume and/or of the number of γH2A.X foci were considered outliers and excluded from the analysis. These two parameters were used to identify outliers because they showed the highest variability among the measured parameters. Data were normalized to the mean of the NT DMSO group for each experimental replicate.

#### CM area coverage in tissue sections

Individual CMs were segmented on MIP images from 4×1µm z-stacks slices (total 4µm), using dedicated Arivis pipeline with watershed (membrane) algorithm on the WGA signal. The sum area of all segmented CMs in a single image was divided by the total image area to calculate CM area coverage.

#### CM nuclei and perinuclear cytoskeleton in tissue sections

Individual CM segments from CM area coverage analysis were superimposed with Hoechst channel to identify and select only CMs with fully covered nuclei and generate a mask in the Hoechst channel with only the selected CMs. The masked Hoechst channel was further background corrected and Otsu algorithm applied to automatically segment the CM nuclei. Further, PN ring segments were created from 0.5µm dilation of the nuclear segments and subtraction of the nucleus. Similarly, cytoplasmic ring segments were created from 3µm dilatation of the PN segments. Nuclear morphology parameters were extracted from the nuclear segments, and mean fluorescence desmin and α-tubulin intensities from the PN and Cyt. segments.

### Statistics

Statistical analysis was performed using Matlab, OriginPro (Version 9 and 2018), Prism (v10.1.2) and R (v.4.0.2). Statistical test and information on biological and technical replicates can be found in the figure legends. When applicable individual nuclei or cells are indicated with open circles with statistical analysis performed on the pooled data. In addition, mean values of individual animal replicates are superimposed and labeled as closed triangles, and connected by lines between the corresponding experimental conditions, for box and bar plots, the mean line is shown, with whiskers denoting standard error (SE) or the standard devition (SD) from the mean as indicated in each figure legend. Statistical tests for each comparison are denoted in the figure legends. Survival curves were generated using the Kaplan–Meier method and differences in survival tested with a Breslow test.

### Computational model

The adult cardiomyocyte is rod shaped and often binucleated, with cylindrical symmetry that is largely preserved with *Lmna* mutations and LINC complex perturbations. For simplicity, we leverage this symmetry and consider an axisymmetric model covering only a quarter of the cardiomyocyte (see Fig. 7A and Table S1). Using the COMSOL multi-physics software, we simulate nuclear morphological changes due to the mutation and LINC complex disruption. Fig. 7A illustrates a typical finite element mesh used for these simulations. According to this figure, the model comprises three main parts: 1) myofibrils (cytoplasm), 2) nucleus, and 3) perinuclear MT cage, which are explained in the following sections.

#### Myofibrils

Adult cardiomyocytes are predominantly filled with myofibrils, encapsulated by the surrounding sarcolemma. Myofibrils are long contractile fibers that are comprised primarily of interdigitating actin and myosin filaments which are precisely held together by titin proteins^56^. Experimental observations show that during diastole, actomyosin interaction is not fully off and thus there is an active contraction in resting cardiomyocytes ^34^. This contraction encounters resistance from restoring forces mediated by titin protein ^34^ and geometric constraints imposed by the myocardium microenvironment ^33^. Our simulations (discussed later) demonstrate that this diastolic contraction laterally compresses central nuclei, causing them to elongate in the axial direction. This nuclear elongation, in turn, compresses the surrounding MT cage, generating pushing elastic forces in the perinuclear MT cage. Consequently, a complex prestress field exists in the cardiomyocyte that governs the nuclear morphology changes. The existence of this resting stress field is supported by the observation that the elongated cardiomyocyte nuclei become round upon isolation ^32^. To induce this stress field in our model, we note that available experiments ^57^ show that myofibrils tend to assemble in the direction of maximum principal stress ^37^. This implies that the myofibrillar organization during maturation is mainly controlled by stress-activated signaling pathways (like calcium pathway) ^33^. Therefore, we employ our previously developed chemo-mechanical model ^36^ to capture this stress dependent assembly of the myofibrils. To this end, we first hypothesize an (imaginary) stress-free configuration for the cardiomyocyte, wherein the nuclei are round, and actomyosin fibers (myofibrils) are randomly distributed and not yet assembled (Fig. 7A). We further assume the following relation between the myofibrillar prestress field (*σ^mf^*), the rest contractility (shown by tensor *ρ_ij_*), and the corresponding strain field (*ε_ij_*) (see ^36^ for more details):

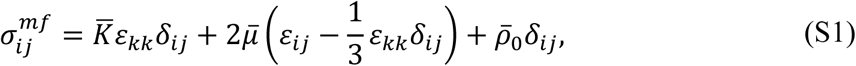

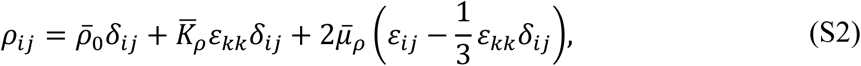

where, we have:

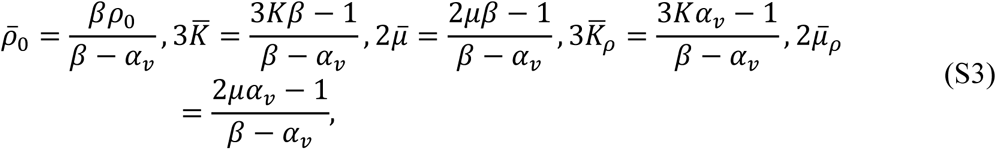

in which β is the chemical stiffness, *α_v_* is the chemo-mechanical feedback parameter, and *K = E*/(3(1 − 2v)) and μ = μ/(2(1 + v)) are the bulk and shear moduli with *E* and v as the elastic modulus and Poisson’s ratio, respectively. Furthermore, ρ_0_is the initial contractility which is zero in the stress-free configuration. The selected values for these model parameters are given in Tables. S2 and S3.

By increasing the initial contractility (i.e., ρ_0_) (Fig. 7B), we then simulate the assembly of myofibrils, resulting in a tendency for the cardiomyocyte to shrink. However, this cell shrinkage is resisted, primarily in the longitudinal direction, by titin proteins and the geometric constraints of the cardiomyocyte microenvironment ^58^, as mentioned earlier. Accordingly, we restrict the longitudinal displacement of the cell at its two ends. Therefore, radial contraction occurs in the cell while its length remains fixed (see Supplementary movie S3). This constrained contraction gives rise to the generation of an anisotropic stress field inside the cytoplasm, as illustrated in Fig. 7C and Extended Data Fig. 6A. As a result of this anisotropic stress field, the contractility tensor *ρ_ij_* will be no longer isotropic (see Eq. S2) since stress activated signaling pathways promote formation of myofibrils in the direction of maximum principal stress ^35,37^. Extended Data Fig. 6B illustrates the obtained directions for the maximum principal stress (maximum tensile stress) field inside the cytoplasm. Notably, there is a great match between these directions and the directions of myofibrils in the physiological conditions.

*<u>Nucleus:</u>* The nucleus is primarily composed of two key mechanical components: chromatin and the nuclear envelope (NE) with its underlying lamina. The NE includes nuclear membranes, and nuclear pore complexes (NPCs), while chromatin serves as the primary building block of the nucleoplasm. Experimental observations ^59^ indicate that chromatin plays a crucial role in resisting small nuclear deformations, while NE and its underlying lamina predominate in scenarios involving large nuclear deformations. These experiments further suggest that the lamina network, especially lamins A and C, exhibits significant strain stiffening, whereas chromatin remains linear even at large deformations. Consequently, we model the nucleoplasm (chromatin) as a linear elastic material with a Young’s modulus of 150 Pa ^35^ and the NE and its underlying lamina as a neo-Hookean hyperelastic layer with an elastic modulus of 15 kPa ^60,61^. This hyperelastic material model captures the strain stiffening of the NE and its underlying lamina upon large deformation. Furthermore, our in vivo observations demonstrate that relative nuclear volume changes due to lamin mutation and LINC complex disruption are less than 10 % (Fig. S6C). Based on these findings, we further consider the nucleoplasm (chromatin) to be nearly incompressible with v = 0.49, while the NE and its underlying lamina are modeled as a fully incompressible hyperelastic layer, similar to the approach in ^60^.

#### Perinuclear MT cage

In adult cardiomyocytes, microtubule organizing centers are predominantly associated with the NE ^62^. MTs anchor to the NE through the LINC complex, particularly nesprin-1 proteins and kinesin motors ^17^, forming a dense and active cage around the nucleus. Consequently, MTs can exert active pushing forces on the nucleus through their polymerization and recruitment of the kinesin motors. This is supported by our super-resolution images (Fig. 6A), which reveal a wavy form of MTs around the nucleus, indicating that these rod-shaped fibers are under compression. To account for these active forces and for simplicity, we introduce an isotropic and homogenous compressive stress field with magnitude *σ_MT_* to the perinuclear MT cage (Eq. S4). We increase this external stress from zero in the stress-free configuration to 500 Pa when inducing the prestress field under WT (physiological) conditions (Fig. 7B). Furthermore, considering the observed microtubule wavelengths, typically between 2-3 μm (Fig. 6A-B), the compressive force experienced by the MTs is estimated to be 100 *pN* or more ^63^. This force is 10 times larger than the force of MT polymerization and 14-15 times larger than the force applied by kinesin motors ^64^. Consequently, we assume that MTs are also under passive elastic compression in adult cardiomyocytes. Accordingly, we model the MT cage as an (active) linear elastic ellipsoid in the stress-free configuration. The constitutive equation for this cage can be written as:

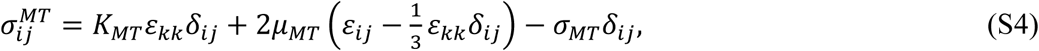

where *K_MT_* and *μ_MT_* are the bulk and shear moduli of the MT cage, respectively. Fig. 7C and Extended Data Fig. 6E show the obtained stress field in the cage. According to these figures, the MT cage imposes maximum compressions on the long and short tips of the nucleus in the longitudinal (z direction) and radial (*r* direction) directions, respectively. This is consistent with our current observations that nuclear ruptures mostly occur at the long tips and also with our recent experimental findings ^3^ that MTs indent into desmin knockdown nuclei mostly around the short sides and in the radial direction.

## Extended Data

**Extended Data Fig. 1:**
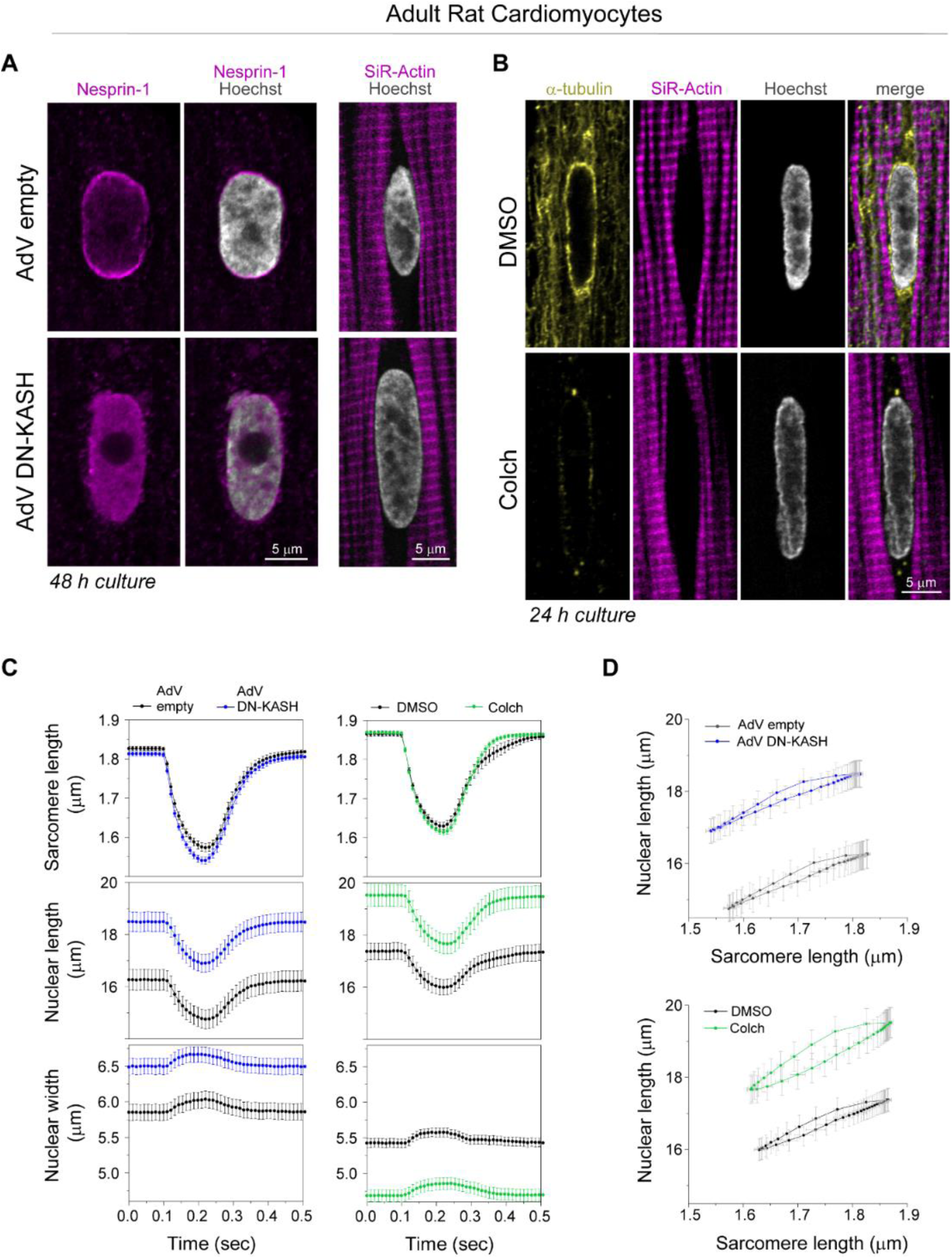
(A) Nesprin-1 perinuclear enrichment in adult rat cardiomyocyte is lost following 48 h of adenoviral DN-KASH transduction, with no apparent changes in perinuclear sarcomeric organization (labeled with SiR-actin). (B) MT network (yellow) is enriched at axial nuclear tips depleted of myofibrils (magenta). Colchicine treatment (1µM) for 24 h results in complete loss of the MT network (including the dense perinuclear MT cage), with no apparent change in perinuclear sarcomeric organization (labeled with SiR-actin). Representative nuclear mid-plane images are shown. (C) Sarcomere length, nuclear length, and nuclear width recordings over time during stimulated cardiomyocyte contraction. Left - cardiomyocytes treated for 48 h in culture with AdV empty (black) or AdV DN-KASH (blue). Right - cardiomyocytes treated for 24 h in culture with DMSO (black) or 1µM colchicine (green). (D) Nuclear length versus sarcomere length coupling plots for AdV DN-KASH and colchicine treatments. AdV empty and AdV DN-KASH (48 h): *N* = 3, *n* = 51. DMSO and colch (24h): *N* = 3, *n* = 51. Data presented as mean ± SE.

**Extended Data Fig. 2:**
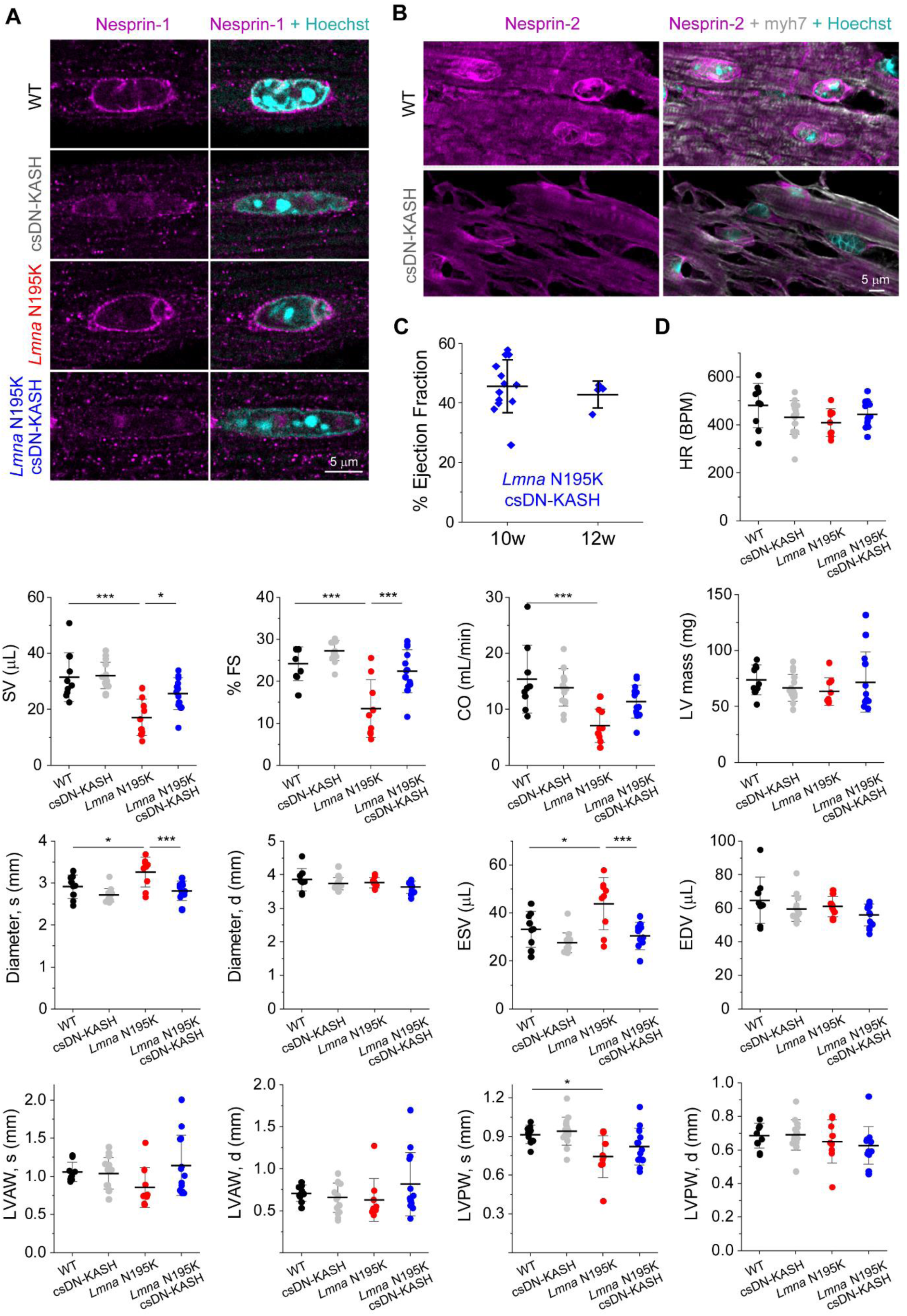
(A) Nesprin-1 perinuclear enrichment in adult WT and *Lmna* N195K mouse cardiomyocytes is lost following cardiac specific in-vivo disruption of the LINC complex (csDN-KASH). Representative nuclear mid-plane images are shown. (B) Nesprin-2 perinuclear enrichment in cardiomyocytes is lost in csDN-KASH mice (top). Representative tissue sections are shown. Cardiomyocyte nuclei identified by myh7 and Hoechst counterstain (bottom) (C) Left ventricular ejection fraction measured by echocardiography, for the *Lmna* N195K csDN-KASH mice at 10 and 12 weeks of age. 10 weeks: *N* = 13, 12 weeks: *N* = 4. Error bars represents mean ± SD. (D) Echocardiography measurements at 10 weeks of age for the indicated groups. Heart rate (HR), stroke volume (SV), % fractional shortening (FS), cardiac output (CO), corrected left-ventricular mass (LV mass), end-systolic and end-diastolic left ventricular diameters (Diameter, s and d), end-systolic and end-diastolic volume (ESV, EDV), end-systolic and end-diastolic left-ventricle anterior and posterior wall thickness (LVAW, s and d, LVPW s and d). WT: *N* = 9, csDN-KASH: *N* = 15, *Lmna* N195K: *N* = 9, *Lmna* N195K csDN-KASH *N* = 13. Error bar represents mean ± SD. Statistical significance determined by one-way ANOVA with Tukey multiple correction. (*, *p* < 0.05, **, *p* < 0.01, ***, *p* < 0.001)

**Extended Data Fig. 3:**
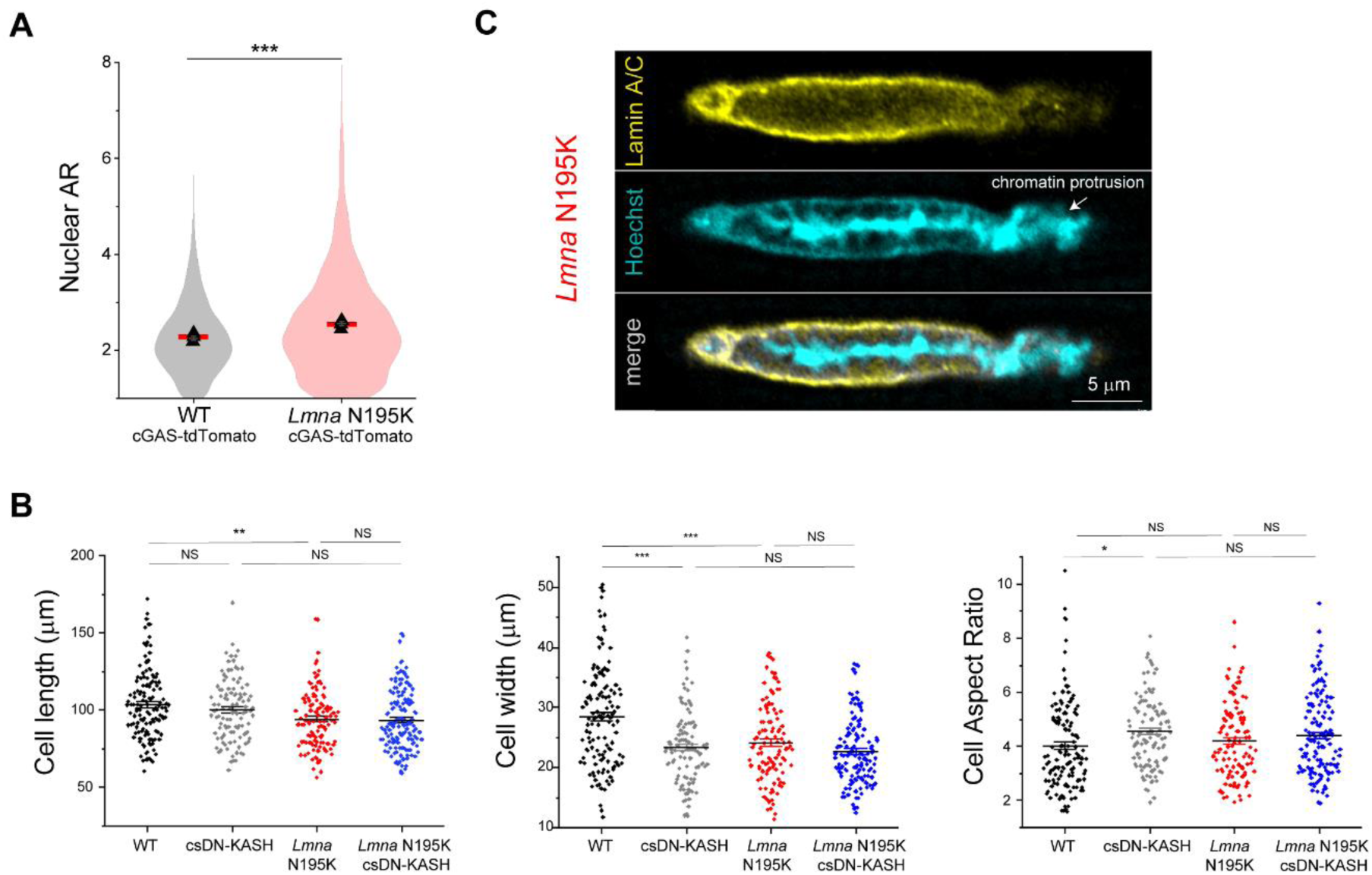
(A) Quantification of nuclear aspect ratio in cardiomyocytes isolated from WT and *Lmna* N195K mice carrying cGAS-tdTomato reporter. Red lines represent mean of pooled nuclei, and black triangles correspond to individual replicate means. WT cGAS-tdTomato: *N* = 4, *n* = 1898. *Lmna* N195K cGAS-tdTomato: *N* = 4, *n* = 2915. (B) Isolated cardiomyocyte cell length, width and cell aspect ratio for the indicated mouse models. WT: *N* = 3, *n* = 133. csDN-KASH: *N* = 3, *n* = 105. *Lmna* N195K: *N* = 3, *n* = 113. *Lmna* N195K csDN-KASH: *N* = 3, *n* = 143. Data presented as mean ± SE. Statistical significance determined by 1-way ANOVA with Bonferroni correction (*, *p* < 0.05, **, *p* < 0.01, ***, *p* < 0.001). (C) NE rupture in *Lmna* N195K cardiomyocytes manifested by chromatin protrusion from the nucleus, with partial lamina coverage. Representative single confocal image of the nucleus is shown.

**Extended Data Fig. 4:**
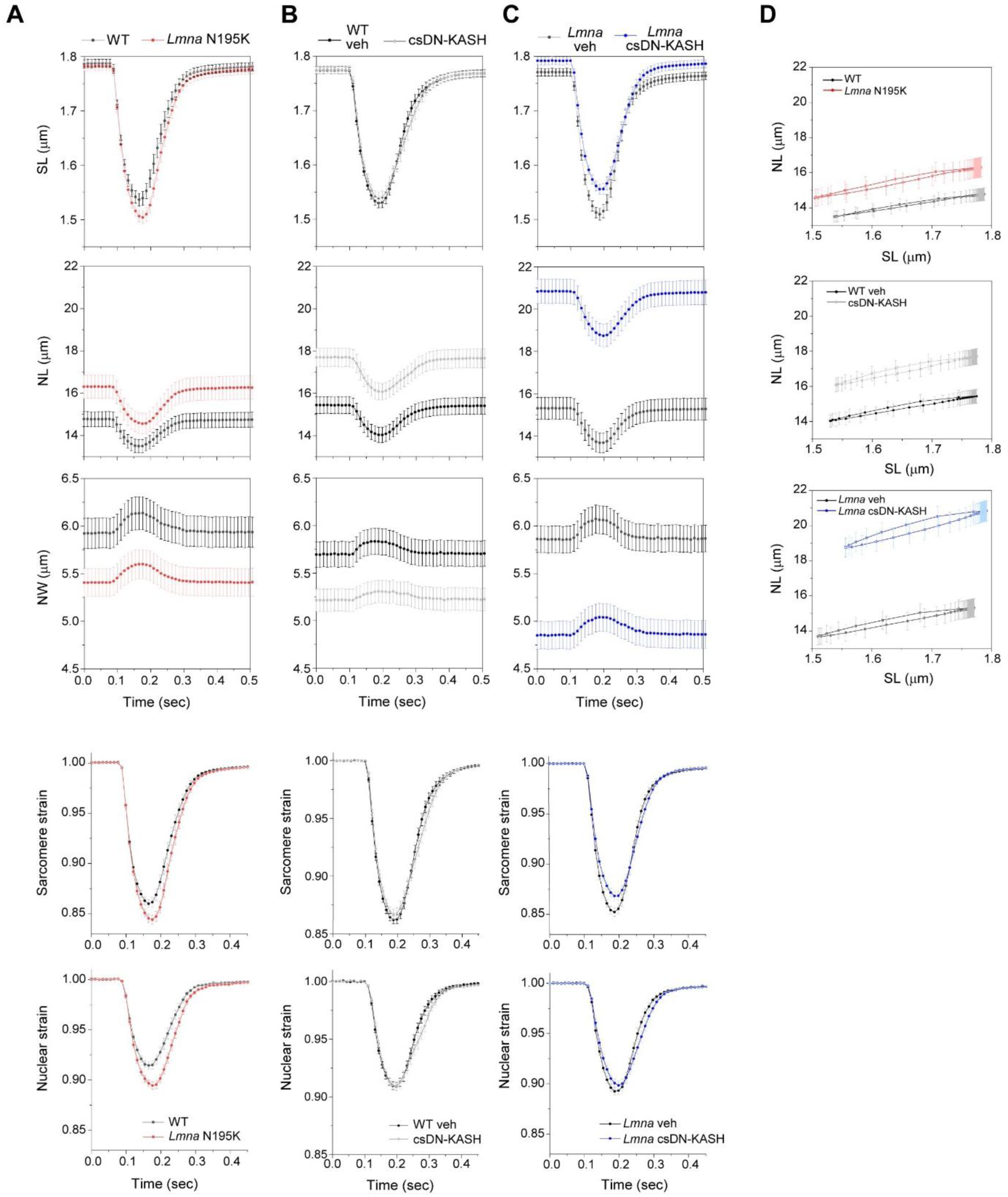
Sarcomere length (SL), nuclear length (NL), and nuclear width (NW) recordings over time during stimulated contraction of freshly isolated cardiomyocytes 8-9 weeks of age. Sarcomere strain and nuclear strain over time are plotted below. (A) *Lmna* N195K (red) and WT littermate controls (black). (B) Cardiac specific LINC complex disruption (gray) and WT vehicle controls (black). (C) *Lmna* N195K with cardiac specific LINC complex disruption (blue) and littermate *Lmna* N195K controls (black). (D) NL versus SL coupling plots for the corresponding experimental groups. WT: *N* = 4, *n* = 58. *Lmna* N195K: *N* = 4, *n* = 69. WT veh: *N* = 4, *n* = 81. csDN-KASH: *N* = 4, *n* = 69. *Lmna* N195K veh: *N* = 4, *n* = 55. *Lmna* N195K csDN-KASH: *N* = 4, *n* = 59. Data presented as mean ± SE.

**Extended Data Fig. 5:**
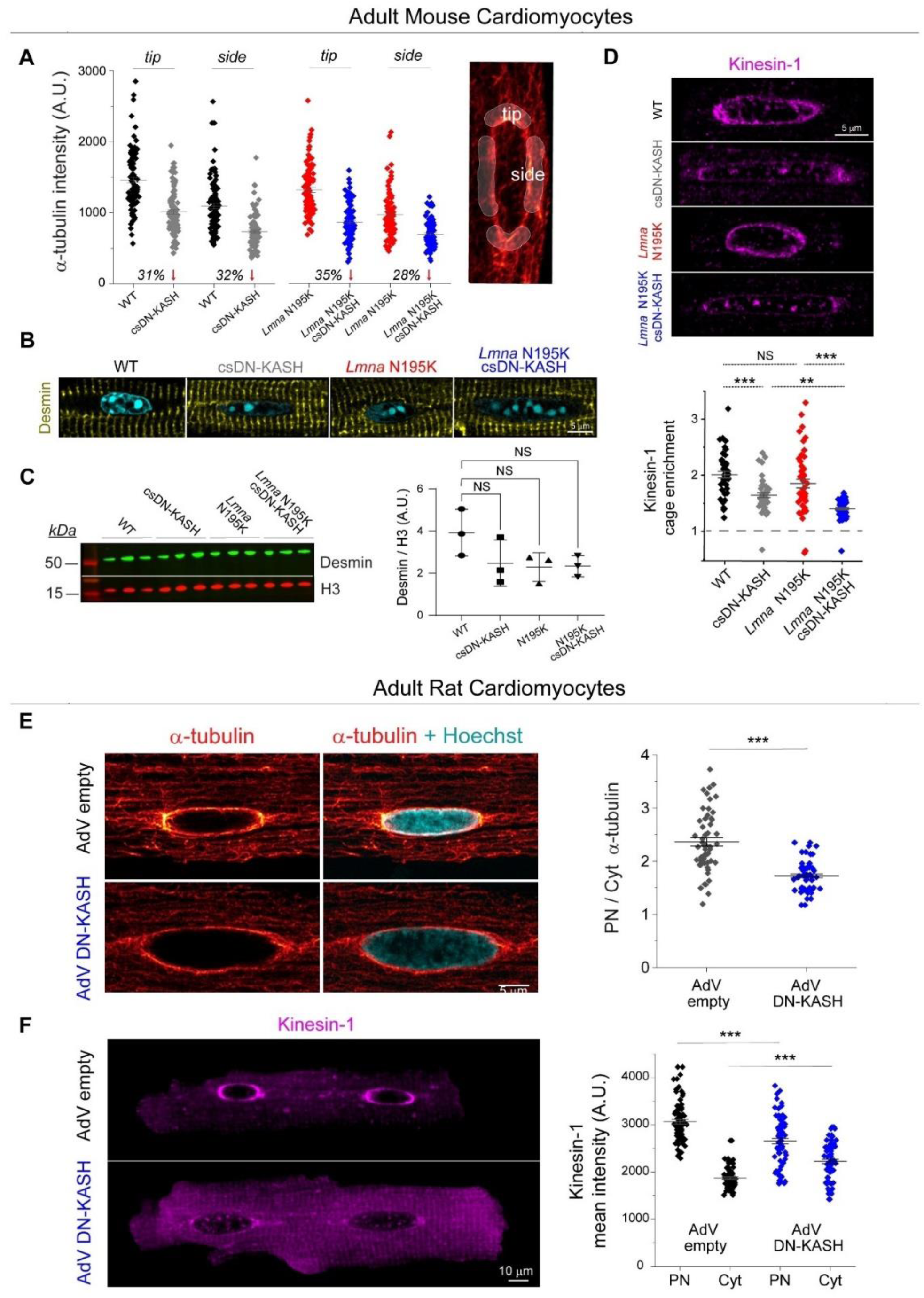
(A) Quantification α-tubulin enrichment at the nuclear tips and sides in the indicated mouse model cardiomyocytes. (B) Representative mid-plane immunofluorescence images of desmin intermediate filament network surrounding the nuclei (Hoechst) of the indicated groups. (C) Desmin western blot and quantification, normalized to H3, for the indicated groups. (D) Representative mid-plane immunofluorescence images of kinesin-1 and quantification of perinuclear kinesin-1 enrichment, defined as PN to Cyt kinesin-1 ratio. WT: *N* = 2, *n* = 46. csDN-KASH: *N* = 2, *n* = 51. *Lmna* N195K: *N* = 2, *n* = 50. *Lmna* N195K csDN-KASH: *N* = 2, *n* = 59. (E) Representative nuclear mid-plane images of adult rat cardiomyocytes labeled with α-tubulin and Hoechst (left), following 48h of adenoviral LINC complex disruption. Quantification of perinuclear α-tubulin enrichment is shown on the right. AdV empty and AdV DN-KASH: *N* = 2, *n* = 29. (F) Representative nuclear mid-plane images of kinesin-1 in the entire adult rat cardiomyocyte following 48h of adenoviral LINC complex disruption (left). Quantification of kinesin-1 intensity separately at the perinucleus and the cytoplasm is shown on the right. AdV empty: *N* = 2, *n* = 78, AdV DN-KASH: *N* = 2, *n* = 80. Data presented as mean ± SE. Statistical significance determined by two-tailed t-test (*, *p* < 0.05, **, *p* < 0.01, ***, *p* < 0.001).

**Extended Data Fig. 6:**
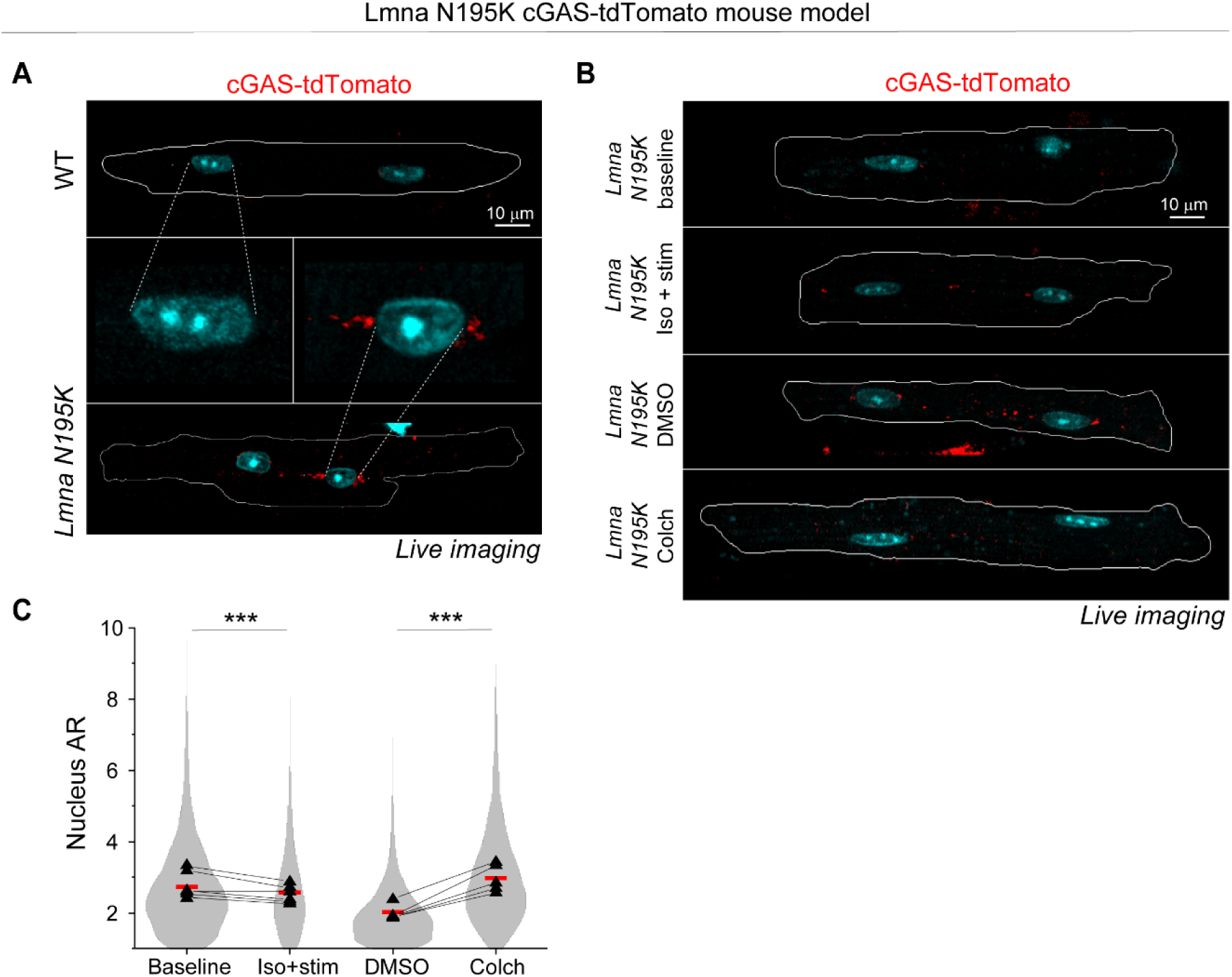
(A) Full cell views of representative maximum intensity projection images labeled for nuclei (Hoechst) and cGAS-tdTomato (red), from freshly isolated cGAS-tdTomato WT, and cGAS-tdTomato *Lmna* N195K mouse models. The middle panel shows a zoom in on the nucleus and the perinuclear region used for cGAS foci quantification. (B) (C) Full cell views of representative maximum intensity projection images labeled for nuclei (Hoechst) and cGAS-tdTomato (red) for the indicated experimental groups. (D) Nuclear aspect ratio (AR) for the experimental groups of cGAS-tdTomato *Lmna* N195K model. *N* = 6, *n* = 4869 (baseline), *n* = 2286 (1 h iso + stim), *n* = 4820 (24 h DMSO), *n* = 4263 (24 h colch). Red lines represent the puled nuclei mean for each experimental group. Superimposed black triangles represent the replicate means and connected by lines to their respective treatment conditions. Statistical significance determined by 1-way ANOVA with Bonferroni correction (*, *p* < 0.05, **, *p* < 0.01, ***, *p* < 0.001).

**Extended Data Fig. 7:**
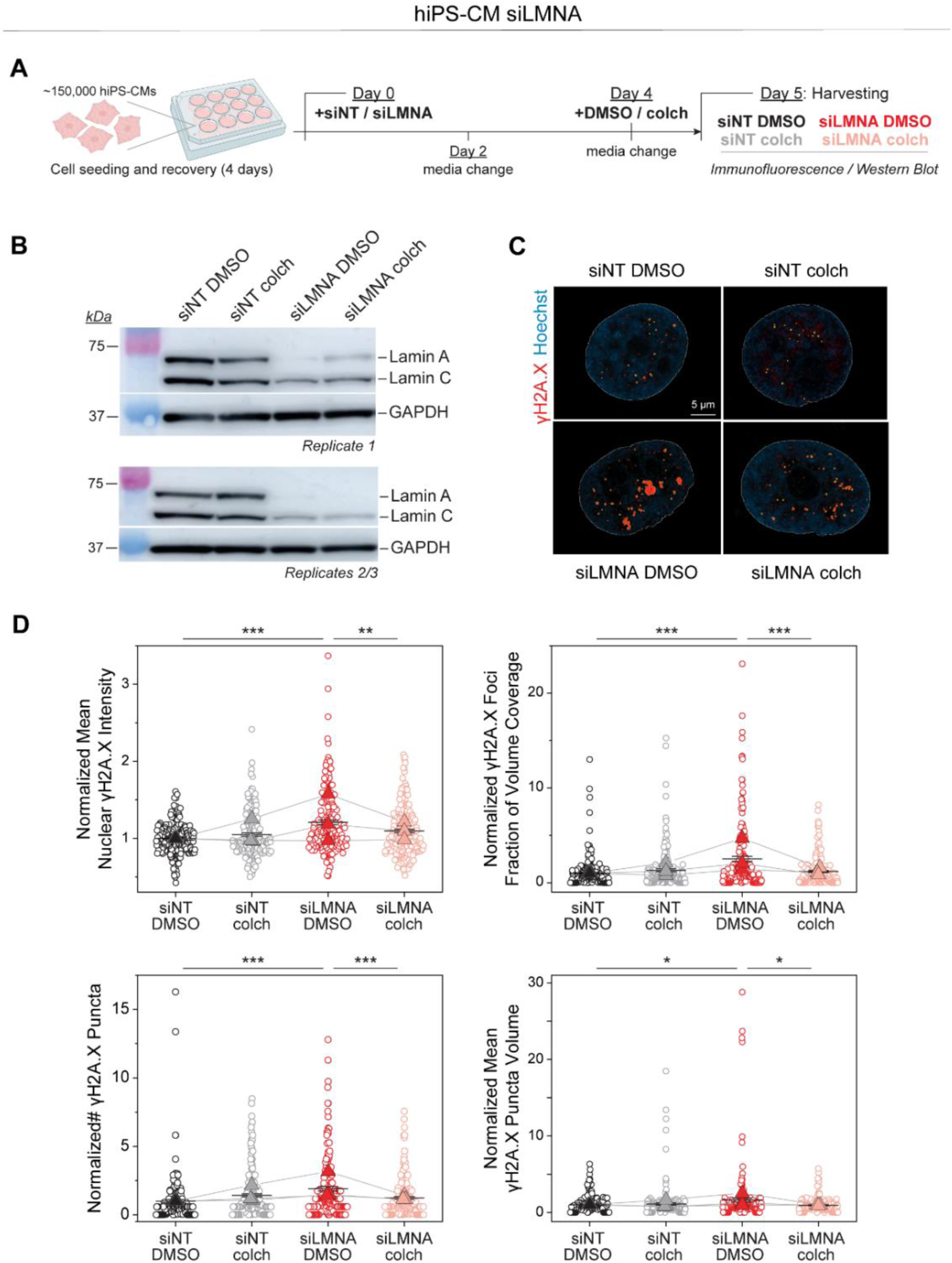
(A) Experimental design for the examination of DNA damage in hiPSC-derived cardiomyocytes upon transfection with non-targeted siRNA (siNT) or LMNA-targeted siRNA (siLMNA), and subsequent DMSO or colchicine (colch) treatment. (B) Western blots, separate for replicate 1 and pooled lysates from replicates 2/3 showing the siRNA-mediated lamin A/C knock-down in hiPSC-derived cardiomyocytes. GAPDH was used as a loading control. (C) Representative mid-plane images of nuclei (Hoechst) and γH2A.X foci, from the four experimental groups of hiPSC-derived cardiomyocytes. Hoechst was used to outline the nuclei (white line), and the detected γH2AX foci are highlighted in yellow. (D) Quantification of mean nuclear γH2A.X intensity and γH2A.X foci fraction of nuclear volume coverage (top). Quantification of the normalized number and volume of γH2A.X foci (bottom). Each replicate is normalized to the mean value of the NT DMSO group. Individual nuclei presented as open circles with mean ± SE of the pooled nuclei. Closed triangles depict replicate means connected by lines to their respective experimental conditions. *N* = 3, *n* = 199 (NT DMSO), *N* = 3, *n* = 197 (NT colch), *N* = 3, *n* = 192 (siLMNA DMSO), *N* = 3, *n* = 203 (siLMNA colch). Statistical significance determined by one-way ANOVA with Bonferroni correction (*, *p* < 0.05, **, *p* < 0.01, ***, *p* < 0.001).

**Extended Data Fig. 8:**
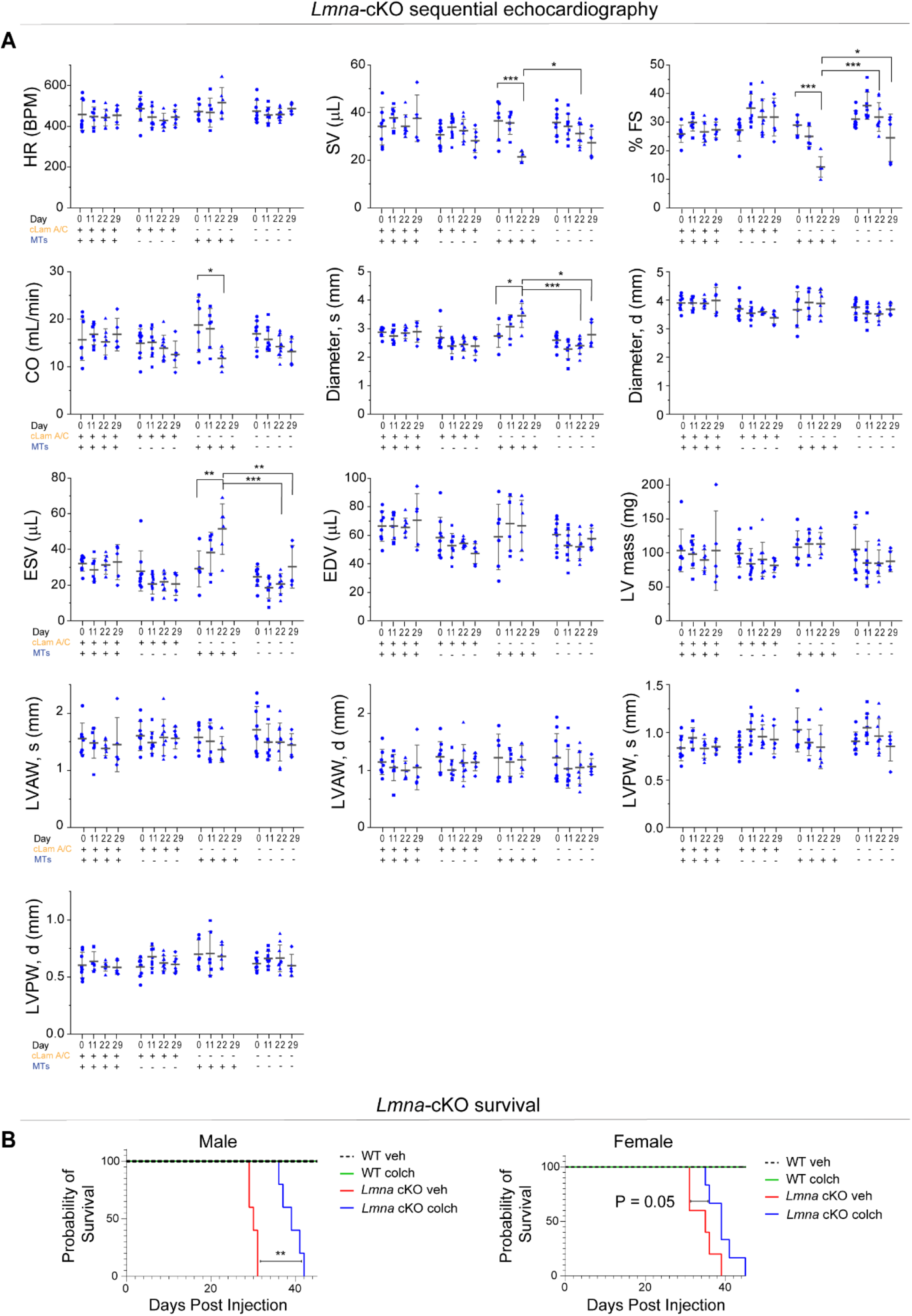
Sequential echocardiography measurements in cardiac specific *Lmna* depleted mice with in-vivo microtubule disruption. Data shown for pre-injection (day 0), 11, 22, and 29 days post initial injection. Herat rate (HR), stroke volume (SV), % fractional shortening (FS), cardiac output (CO), end-systolic and end-diastolic left ventricular diameters (Diameter, s and d), end-systolic and end-diastolic volume (ESV, EDV), corrected left-ventricular mass (LV mass), end-systolic and end-diastolic left-ventricle anterior and posterior wall thickness (LVAW, s and d, LVPW s and d). WT + veh: *N* = 8 (day 29: *N* = 5), WT + colch: *N* = 9 (day 29: *N* = 6), *Lmna* cKO + veh: *N* = 6 (*Lmna* cKO + vehicle treated mice do not survive to day 29 post injection), *Lmna* cKO + colch: *N* = 9 (day 29: *N* = 5). Error bars represent mean ± 1 SD. B) Kaplan-Meier survival plots of the different experimental groups. Male: WT+veh: *N* = 2, WT+colch: *N* = 2, *Lmna* cKO+veh: *N* = 5, *Lmna* cKO+colch: *N* = 5. Female: WT+veh: *N* = 3, WT+colch: *N* = 2, *Lmna* cKO+veh: *N* = 5, *Lmna* cKO+colch: *N* = 6. Statistical significance determined by two-way ANOVA with Tukey multiple correction. (*, *p* < 0.05, **, *p* < 0.01, ***, *p* < 0.001)

**Extended Data Fig. 9:**
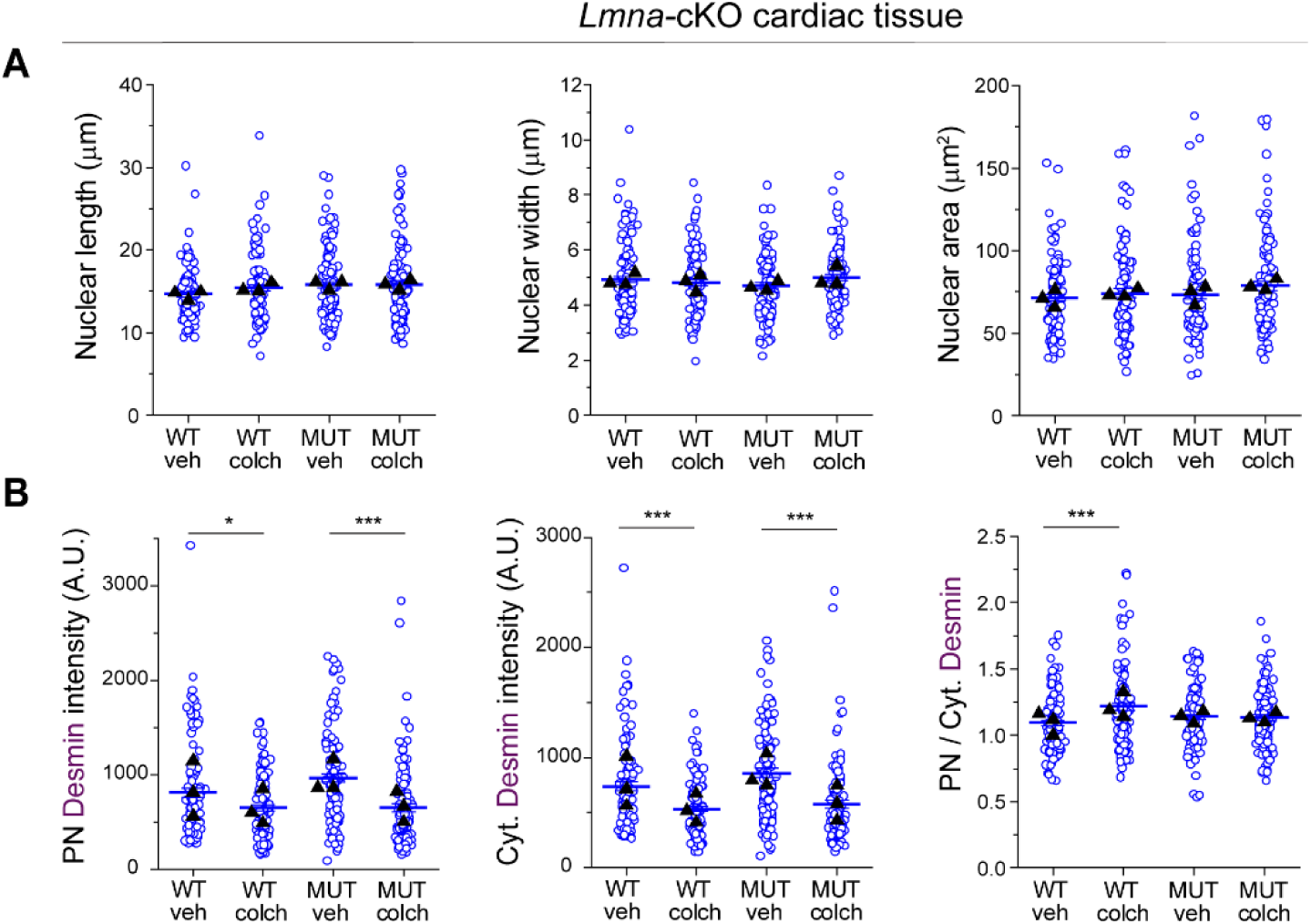
(A) Quantification of cardiomyocyte nuclear length, width, and area. (B) Quantification of perinuclear (PN), cytoplasmic (Cyt) and PN/Cyt. desmin intensity from maximum-intensity-projection images of cardiac tissue sections for the indicated groups. *N*=3 mice per group. WT veh: *n* = 113. WT colch: *n* = 112. *Lmna* cKO (MUT) + veh: *n* = 111. *Lmna* cKO + colch: *n* = 112 nuclei. Statistical significance determined by 1-way ANOVA with Bonferroni correction. (*, *p* < 0.05, **, *p* < 0.01, ***, *p* < 0.001)

**Extended Data Fig. 10:**
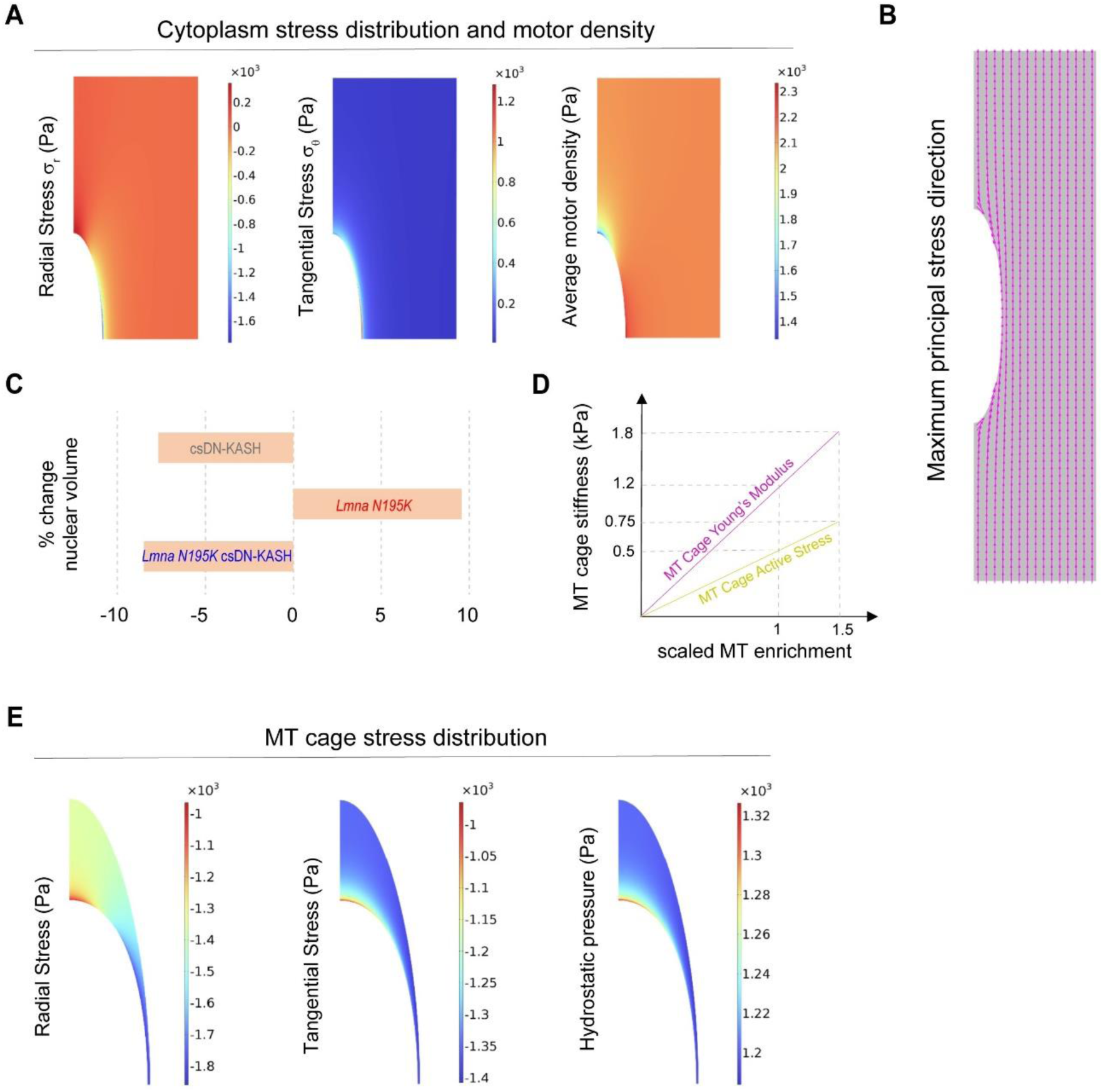
(A) Simulation results depicting stress distribution and average myosin motor density in the cardiomyocyte cytoplasm under physiological conditions. Radial stress component, Tangential stress component and average motor density defined as (ρ_11_ + ρ_22_ + ρ_33_)⁄3. (B) Model predictions for the maximum principal (maximum tensile) stress direction in the cytoplasm. (C) In vivo % volume changes relative to WT conditions. (D) Variations of the MT cage stiffness and active stress to simulate MT enrichment around the nucleus. (E) Simulation results for stress distribution in the microtubule cage in WT conditions. Radial stress component (Pa), Tangential stress component (Pa) and hydrostatic pressure (Pa).

---

**Table S1:** Geometry parameters of the model parts in the stress-free configuration.

| Parameter | value |
| --- | --- |
| Cell radius | 35 $\mu m$ |
| Cell length | 100 $\mu m$ |
| Nuclear radius | 4 $\mu m$ |
| NE + lamina thickness | 0.2 $\mu m$ |
| MT cage long axis length | 8 $\mu m$ |
| MT cage short axis length | 4.4 $\mu m$ |

**Table S2:**
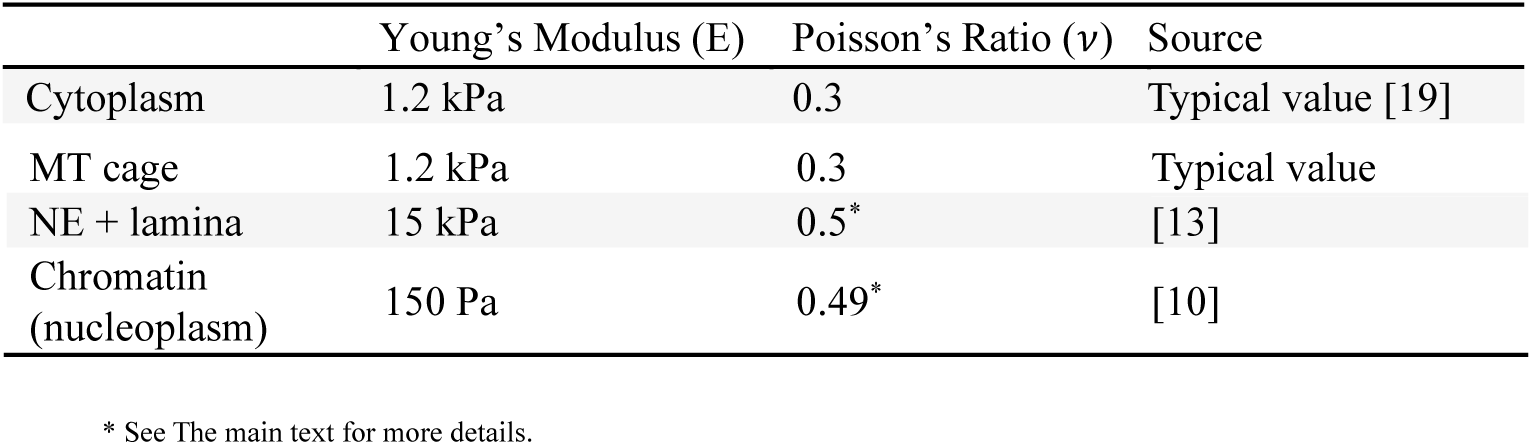
Material properties of the model parts.

**Table S3:**
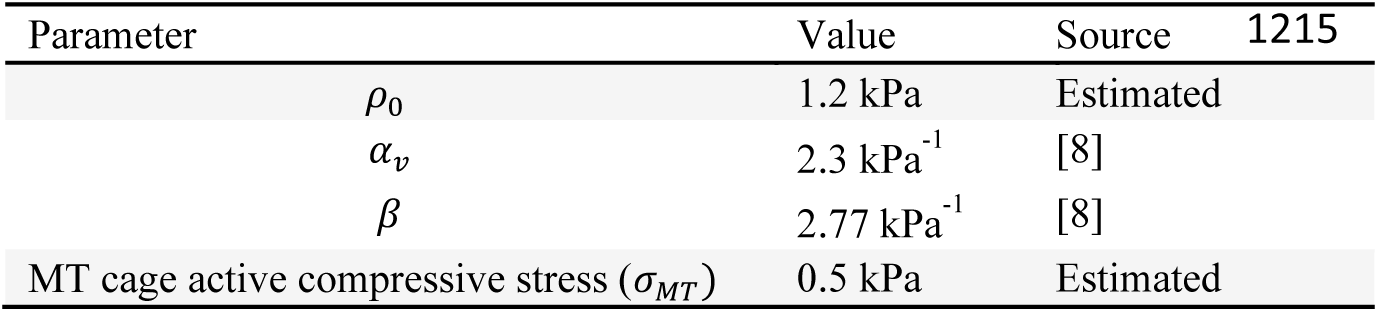
Parameters for the active deformations of the cytoplasm and the MT cage.

## Supplementary Movies

**Supplementary Movie S1:** Electrically stimulated (1 Hz) adult rat cardiomyocyte labeled with SiR-actin and Hoechst to visualize sarcomeres and DNA, respectively.

**Supplementary Movie S2:** Electrically stimulated (1 Hz) adult rat cardiomyocyte labeled with SPY-555 tubulin and Hoechst demonstrating MT cage buckling during contraction.

**Supplementary Movie S3:** Cardiomyocyte model simulation following assembly of myofibrils (increased myofibril prestress with constant cytoplasm length) demonstrating development of anisotropic stress field within the cell with overall cellular radial compression and nuclear elongation.

## Notes

### Competing Interest Statement

The authors have declared no competing interest.

### Summary of Updates

Additional cardiac function and structure data in Figure 3. Also new Figure 7 now includes extended data for in-vivo cardiac specific Lmna depletion mouse model with survivability analysis, cardiac structure and function, and cellular characterization of the microtubule network and nuclear integrity. The new data further demonstrates that in-vivo disruption of the microtubule network protects from nuclear ruptures related to Lmna depletion in cardiomyocytes.

